# Melanoma-secreted Amyloid Beta Suppresses Neuroinflammation and Promotes Brain Metastasis

**DOI:** 10.1101/854885

**Authors:** Kevin Kleffman, Grace Levinson, Eitan Wong, Francisco Galán-Echevarría, Richard Von-Itter, Indigo Rose, Lili Blumenberg, Alfredo Floristán, James Tranos, Diana Argibay, Jenny Chen, Avantika Dhabaria, Eleazar de Miera Sainz de Vega, Robert Rogers, Youssef Zaim-Wadghiri, Paul Mathews, Iman Osman, Kelly Ruggles, Beatrix Ueberheide, Shane A. Liddelow, Ronald DeMattos, Yue Ming Li, Robert J. Schneider, Eva Hernando

**Author notes:** **Corresponding Author** Correspondence should be addressed to Eva Hernando.

## Abstract

Brain metastasis is a significant cause of morbidity and mortality in multiple cancer types and represents an unmet clinical need. The mechanisms that mediate metastatic cancer growth in the brain parenchyma are largely unknown. Melanoma, which has the highest rate of brain metastasis among common cancer types, is an ideal model to study how cancer cells adapt to the brain parenchyma. We performed unbiased proteomics analysis of melanoma short-term cultures, a novel model for the study of brain metastasis. Intriguingly, we found that proteins implicated in neurodegenerative pathologies are differentially expressed in melanoma cells explanted from brain metastases compared to those derived from extracranial metastases. This raised the exciting hypothesis that molecular pathways implicated in neurodegenerative disorders are critical for metastatic adaptation to the brain.

Here, we show that melanoma cells require amyloid beta (Aβ), a polypeptide heavily implicated in Alzheimer’s disease, for growth and survival in the brain parenchyma. Melanoma cells produce and secrete Aβ, which activates surrounding astrocytes to a pro-metastatic, anti-inflammatory phenotype. Furthermore, we show that pharmacological inhibition of Aβ decreases brain metastatic burden.

Our results reveal a mechanistic connection between brain metastasis and Alzheimer’s disease – two previously unrelated pathologies, establish Aβ as a promising therapeutic target for brain metastasis, and demonstrate suppression of neuroinflammation as a critical feature of metastatic adaptation to the brain parenchyma.

## Main

Brain metastasis is the most common form of adult intracranial malignancy^1^ and results in severe morbidity and mortality. 40-75% of Stage IV melanoma patients develop brain metastasis^2, 3^, reflecting melanoma’s striking ability to colonize the brain. Brain metastases are less responsive than extracranial metastases to current cancer therapies^4–6^, and the majority of patients succumb to disease in less than one year^7^. Furthermore, patients with brain metastasis are often excluded from clinical trials and urgently need new clinical options. In recent years, research has started to elucidate the molecular mechanisms contributing to the multi-step process of brain metastasis. Most findings have focused on cancer extravasation across the blood-brain barrier (BBB), which cannot be leveraged therapeutically given that the vast majority of brain metastasis patients will present with extravasated cancer cells at the time of cancer diagnosis. The main bottleneck in the brain metastatic process has been shown to be the successful expansion of a single cell in the brain parenchyma to form a macro-metastasis^8^. Recent studies have begun to demonstrate the role of the brain microenvironment in this process. In particular, reactive astrocytes have been shown to interact with cancer cells in the brain^9, 10^ and exhibit both pro- and anti-brain metastatic activity^11, 12^. Astrocytes have several roles in normal brain physiology, including neurotransmitter uptake^13^, metabolic support^14^, and response to injury^15^. Furthermore, astrocytes have been heavily implicated as both neurotoxic and neuroprotective in a variety of neurodegenerative pathologies, including Alzheimer’s disease^16–18^. These findings suggest the intriguing possibility of a functional connection between neurodegenerative pathologies and brain metastasis, which has not yet been explored.

Here, we demonstrate that melanoma cells require amyloid beta (Aβ), a polypeptide heavily implicated in Alzheimer’s disease, for survival and late growth in the brain parenchyma. Melanoma cells cleave Amyloid Precursor Protein (APP) to produce and secrete Aβ, and Aβ secreted from cancer cells triggers local astrocytes to adopt a pro-metastatic, anti-inflammatory phenotype. Targeting Aβ production by pharmacologic inhibition of *β*-secretase activity suppresses metastatic growth of human cancer cells in the brain parenchyma of mice.

### Proteomics analysis links melanoma brain metastasis and neurodegeneration

To study mechanisms of brain metastasis, we leveraged pairs of brain metastasis-derived (BM) and non-brain metastasis-derived (NBM) melanoma short term cultures (STCs) obtained from the same patient (Figure 1a) as a novel model of brain metastasis. Comparing patient-matched BM and NBM STC pairs reduces the confounding inter-patient heterogeneity characteristic of melanoma. Upon intracardiac injection in immunocompromised mice, BM STCs exhibited an increased ability to metastasize to the brain than their paired NBM STCs (Figure 1b-d, Extended Data Figure 1a,b). Therefore, any molecular differences between paired BM and NBM STCs can be associated with differential brain metastatic ability.

**Figure 1:**
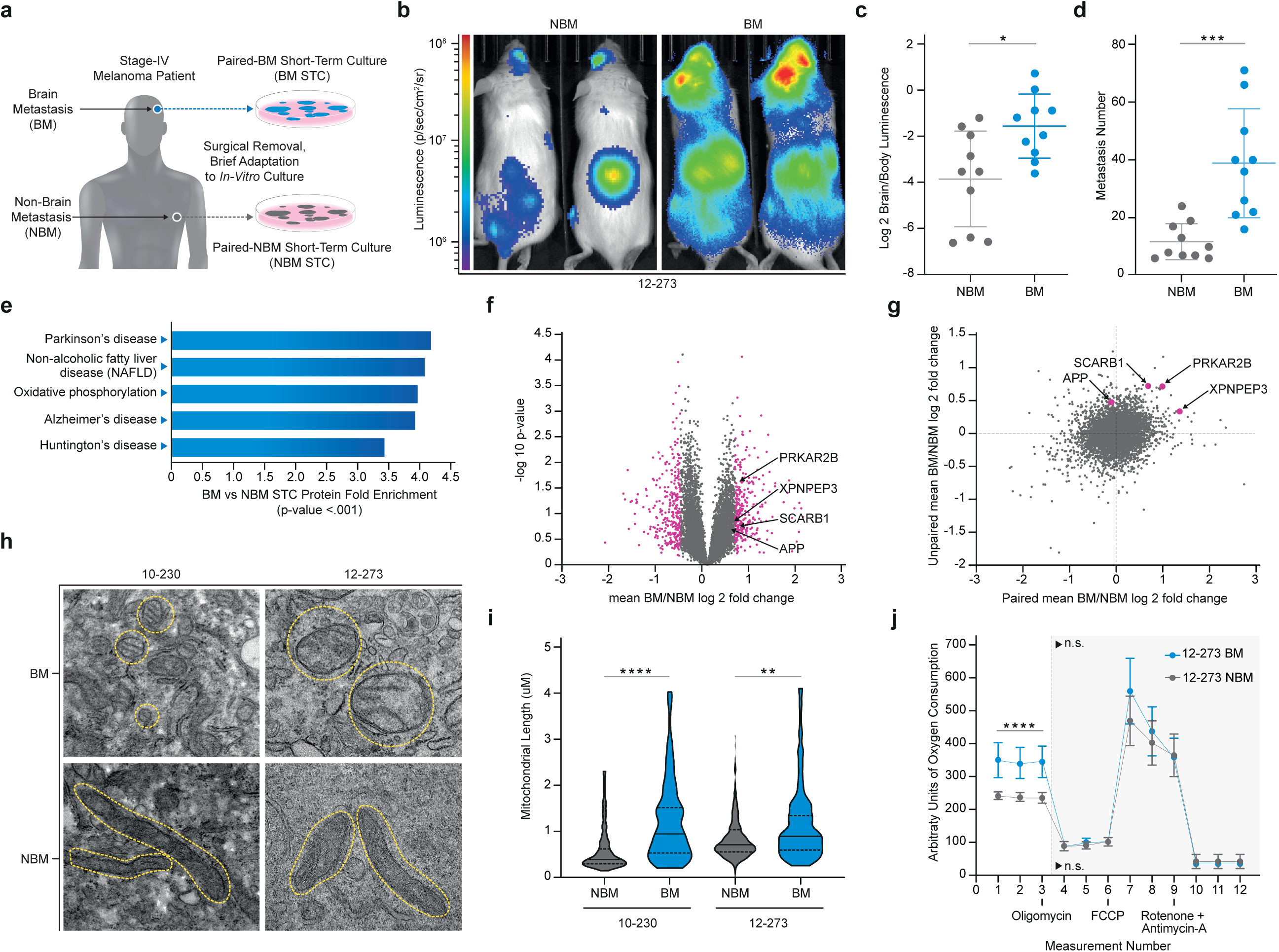
Proteomics Links Melanoma Brain Metastasis and Neurodegeneration. **a**, Diagram of the generation of patient-matched STC pairs. **b**, Representative IVIS images of 12-273 STC pair at 29 days post-intracardiac injection in mice. **c**, Quantified brain/body luminescence ratio on day 29 on 12-273 NBM vs BM (* p<0.05). **d**, Number of brain metastases quantified by microscopy of serial FFPE sections with H&E staining. (12-273 NBM vs BM, p<0.0005; n=2 independent experiments with 10 mice per group, representative experiment shown). **e-g**, Mass spectrometry analysis of whole cell lysates from a cohort of 14 BM and 11 NBM STCs, including 3 patient-matched pairs. **e**, Top enriched pathways (p<0.001) in BM vs NBM STCs identified from KEGG Pathway analysis of global protein levels. **f**, Volcano plot of mean Log2 BM/NBM fold change of global protein levels and -Log10 p-values. Pink– proteins with mean Log2 BM/NBM fold change >0.6 (1.5 BM/NBM fold change) or < −0.6 (2/3 BM/NBM fold change). **g**, Comparison of mean paired STC BM/NBM Log2 fold change to unpaired STC BM/NBM mean Log2 fold change. Pink – candidates selected for in-vivo mini-screen. **h**, Representative electron microscopy images of paired STCs. Yellow circles outline mitochondria. **i**, Quantification of average mitochondrial length in paired STCs. 10-230 NBM vs BM (**** p<.00005), 12-273 NBM vs BM (** p<.005). **j**, Seahorse MitoStress analysis of oxygen consumption rate in 12-273 NBM and BM. Basal oxygen consumption rate of 12-273 NBM vs BM (**** p<.00005; n=3 independent experiments, 4-6 biological replicates per group per experiment. representative experiment shown)

Using a cohort of 15 BM and 11 NBM derived STCs, which included 3 isogenic pairs, we performed an unbiased mass spectrometry analysis of whole cell protein lysates to identify novel candidate pathways and proteins that may mediate melanoma brain metastasis. KEGG pathway analysis of differentially expressed proteins revealed an enrichment in proteins related to neurodegenerative pathologies and oxidative phosphorylation in the BM vs NBM STCs (Figure 1e-g). Proteomics results were validated by Western blot analyses (Extended data Fig. 1c,d). Metabolic profiling demonstrated that BM STCs have increased mitochondrial fusion and electron density (Figure 1h,i), elevated mitochondrial oxygen consumption (Fig. 1j and Extended data Figure 1e,g,i), and decreased glycolysis (Extended data Figure 1f,h,j) than their respective paired NBM. These results provide further evidence of a recently reported connection between melanoma brain metastasis and oxidative phosphorylation^19, 20^, supporting the capability of our proteomic results to reflect phenotypic differences related to brain metastatic ability.

### APP is specifically required for melanoma brain metastasis

We performed an in vivo brain metastasis mini-screen to identify novel mediators of melanoma brain metastasis. We first selected three proteins (PRKAR2B, SCARB1, XPNPEP3) found consistently increased in both paired and unpaired BM STCs relative to NBM STCs (Figure 1g), and functionally related to neurodegeneration and/or mitochondrial metabolism. Additionally, a review of the literature of top differentially expressed proteins from the proteomics results revealed several connections with Alzheimer’s disease, APP cleavage, and Aβ. Although APP was not identified as significantly increased in BM vs NBM STCs, proteomics analysis of whole cell lysates cannot detect secreted proteins such as Aβ cleaved from APP. Given that APP and its cleavage products can have profound effects on the brain microenvironment in the development of Alzheimer’s disease ^21^, we hypothesized that melanoma cells may require APP and/or its cleavage products for survival or growth in the brain parenchyma.

Using lentiviral shRNA, we silenced the four selected candidates in a STC, 12-273 BM, and measured effects on in vitro proliferation (Figure 2b, Extended data fig. 2a-d) and metastatic capability upon intracardiac injection in mice. Silencing of APP and PRKAR2B resulted in a significantly decreased brain to body luminescence signal (Figure 2a), indicating that these proteins have differential effects on metastatic potential to the brain as compared to other organs. However, since mice injected with PRKAR2B-depleted cells had increased metastasis to extracranial organs (Extended Data Figure 1e-g), we decided to narrow our focus to APP. Silencing of APP did not affect the proliferative capacity of melanoma cells *in-vitro* (Figure 2b), but resulted in reduced colonization of the brain and a decreased brain/body luminescence signal (Figure 2c,d). *Ex-vivo* MRI of mouse brains revealed that loss of APP decreases overall brain tumor burden (Figure 2e,f; Supplementary Video 1), number of brain metastases (Figure 2g), and average brain metastasis size (Fig. 2h). Metastatic burden to specific organs was further quantified by NuMA immunohistochemistry, which specifically labels the nuclei of human cells within mouse organs. Loss of APP resulted in a dramatic reduction of brain metastatic burden (Figure 2i,j), but had no significant effect on metastasis to either the liver (Figure 2k) or the kidneys (Figure 2l). Silencing of APP in a second STC, WM-4265-2 BM^22^, also inhibited brain metastasis without significant effects on metastasis to other organs (Extended Data Figure 2h-j) Thus, APP is required for melanoma to colonize the brain but not to metastasize to other organs.

**Figure 2:**
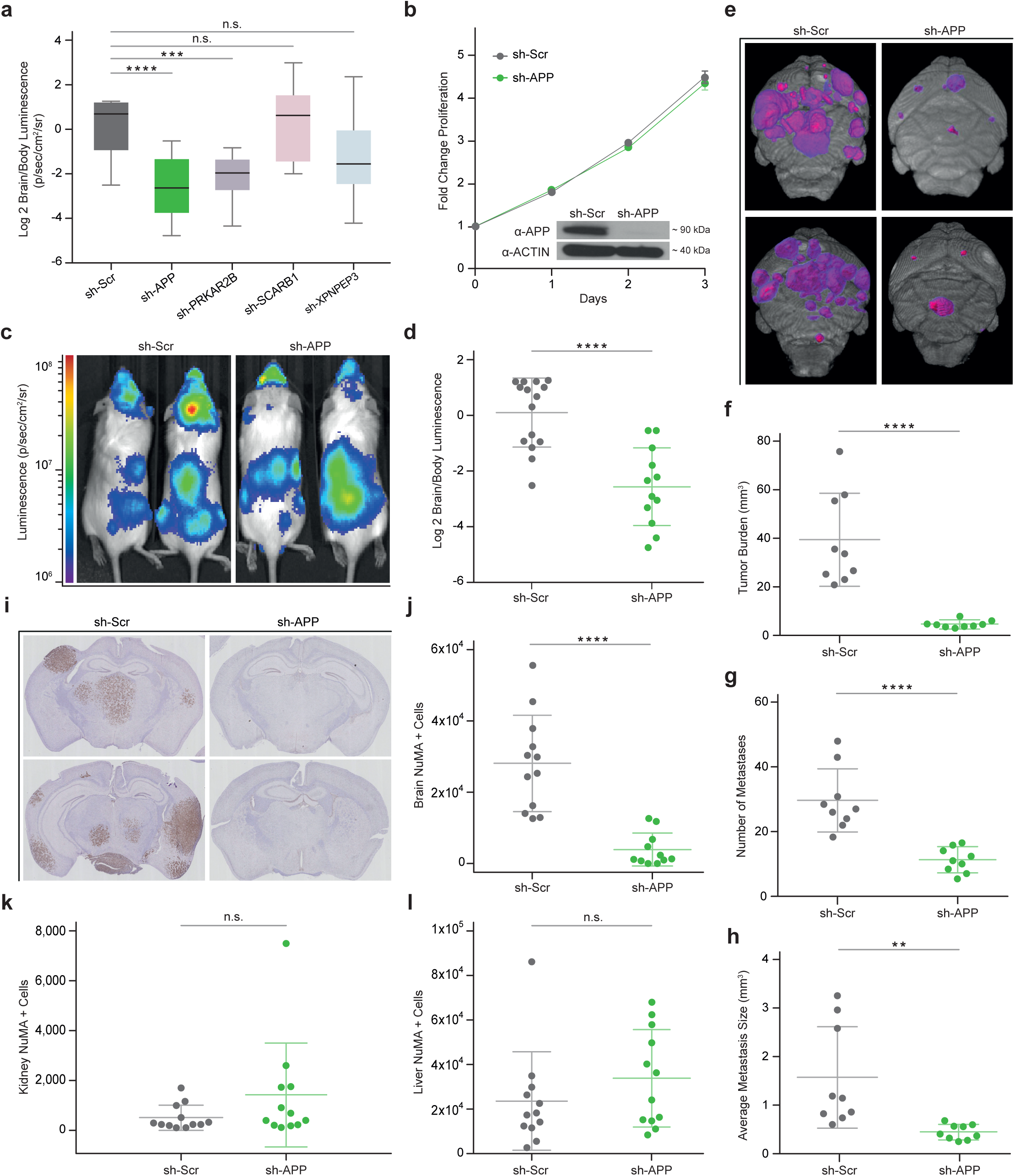
APP is Specifically Required for Melanoma Brain Metastasis. **a**, Quantified Log2 Brain/Body luminescence 35 days post-intracardiac injection in mice of 12-273 BM with shRNA-mediated silencing of selected candidates or scrambled hairpin control (sh-Scr). sh-Scr vs sh-APP (**** p<0.00005), sh-Scr vs sh-PRKAR2B (** p<0.005). (n= 1 experiment, 10-12 mice per group. Box = Interquartile range. Error bars = min to max.) **b**, Fold change in-vitro proliferation and western blot analysis of 12-273BM cells transduced with sh-APP vs sh-Scr. **c**, Representative IVIS images at day 35. **d**, Quantified Log2 Brain/Body luminescence at day 35 of mice injected with 12-273BM cells transduced with sh-Scr vs sh-APP lentivirus (**** p<0.00005). (e-g), Ex-vivo brain MRI of mice injected with 12-273BM sh-Scr vs sh-APP: (n=1 experiment, 9 mice per group). **e**, Representative images. Pink-purple – brain metastasis. Quantification of **f**, brain metastatic burden (**** p<0.00005); **g**, number of brain metastases (**** p<0.00005) and **h**, average metastasis size by MRI. (** p<0.005) (i-l), Labeling of metastatic cells by anti-NuMA immunohistochemistry on FFPE brain slides of mice injected with sh-Scr vs sh-APP: i, Representative brain images; (**j-l**), Quantification of NuMA+ metastatic cells in (**j**) mouse brain (**** p<0.00005), (**k**) kidneys, and (**l**) livers.

### Amyloid Beta is the form of APP required for melanoma brain metastasis

APP can be processed to produce amyloid beta (Aβ), a polypeptide heavily implicated in Alzheimer’s disease, through sequential cleavage by beta and gamma secretases. Although APP is expressed in a wide variety of normal tissues and tumor types, potential Aβ production by cancer cells has never been explored. Therefore, we sought to investigate whether melanoma cells can produce Aβ and if so, whether Aβ production is altered in BM vs NBM STCs. Using probe specific gamma-secretase assays^23^, we established that melanoma cells can cleave APP with gamma secretase and observed consistently increased cleavage of APP in BM vs paired NBM STCs (Figure 3a). Notably, increased cleavage of NOTCH, a canonical gamma secretase substrate, was not consistently observed in paired BM vs NBM STCs (Figure 3b), which suggests a specific association between APP cleavage and brain metastasis. Analysis of melanoma conditioned media demonstrated that melanoma cells are able to produce and secrete Aβ, and that Aβ secretion is increased in BM relative to paired NBM STCs (Figure 3c). Therefore, we posited that Aβ is the specific form of APP required for melanoma brain metastasis.

**Figure 3:**
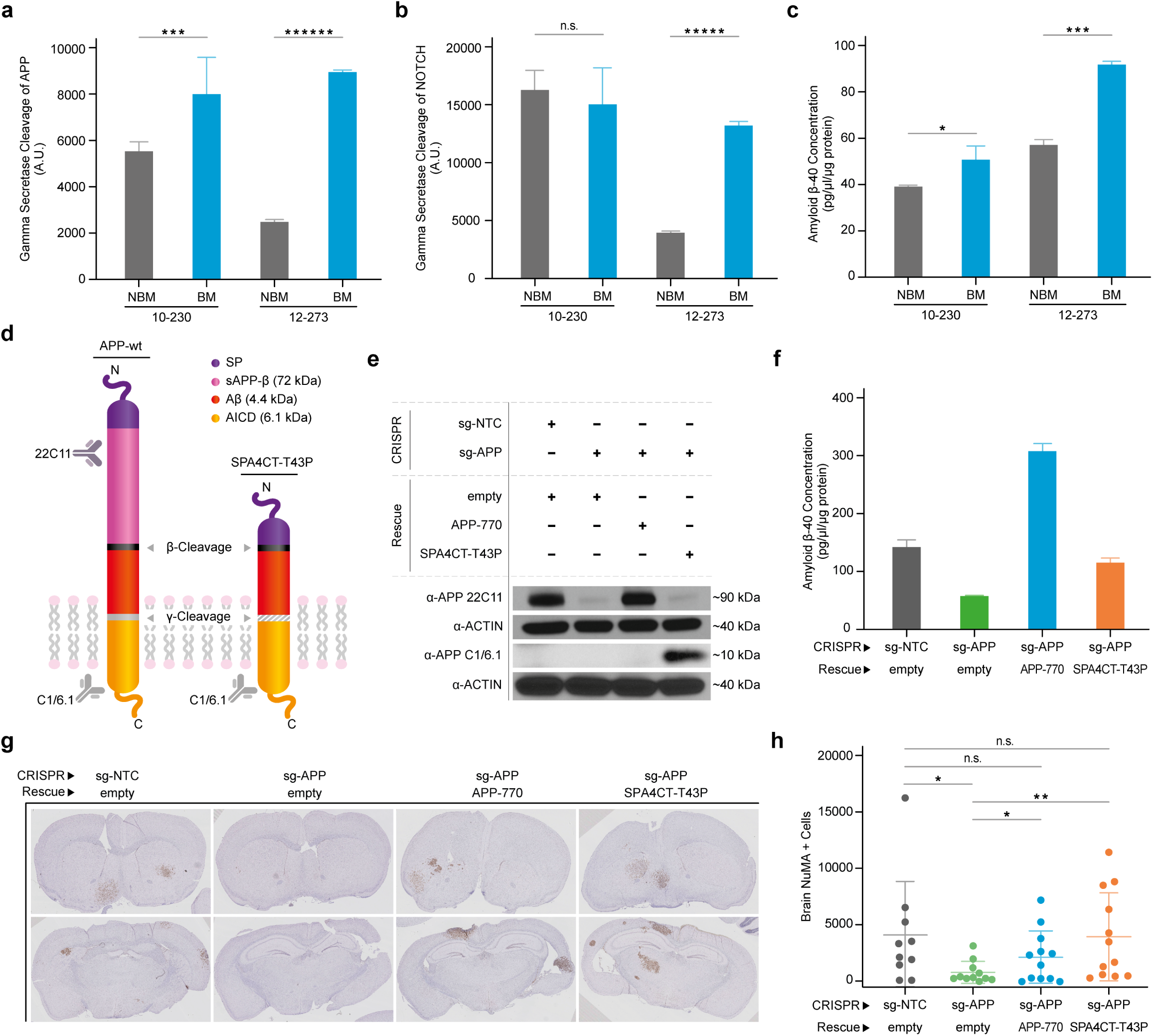
Amyloid Beta (Aβ) is the form of APP required for Melanoma Brain Metastasis. **a**, Quantification of gamma-secretase cleavage of APP. 10-230 NBM vs BM (* p<0.05), 12-273 NBM vs BM (****** p<0.0000005). **b**, Quantification of gamma-secretase cleavage of NOTCH. 12-273 NBM vs BM (***** p<0.000005) (two independent experiments, n=4 biological replicates for group, representative experiment shown). **c**, Quantification of Aβ secretion by ELISA. 10-230 NBM vs BM (* p<0.05), WM-4071 NBM vs BM (*** p<0.0005) (two independent experiments, n=2-4 biological replicates for group, representative experiment shown). **d**, Diagram of wildtype APP and SPA4CT-T43P. **e**, Western blot analysis using anti-APP 22C11 (using actin as loading control) in 12-273 BM infected cells. **f**, Quantification of Aβ secretion by ELISA in 12-273 BM infected cells. **g**, Representative images of FFPE brain slides with labeling of metastatic cells by anti-NuMA immunohistochemistry. **h**, Quantification of NuMA+ metastatic cells in mouse brains. sg-NTC vs sg-APP (* p<0.05), sg-APP vs APP-770 (one sided t-test * p<0.05), sg-APP vs SPA4CT-T43P (one sided t-test ** p<0.005). (n = 10-12 mice per group).

To test this hypothesis, we asked whether Aβ is sufficient to rescue the inhibition of metastasis observed upon APP silencing. We cloned SPA4CT-T43P ^24^, a heavily truncated (>75% of APP amino acid sequence removed), mutant form of APP that retains the ability to produce Aβ but cannot produce other major cleavage products, such as sAPP-*α* and sAPP-*β* (Figure 3d). The substitution of proline for threonine at position 43 partially inhibits gamma secretase cleavage^25^ and prevents excessive Aβ generation (Extended Data Figure 3a,b). We knocked out APP in 12-273 BM STC using CRISPR/Cas9 and reintroduced either APP-770, the major full-length APP isoform expressed in melanoma cells (Extended Data Figure 3c), or SPA4CT-T43P (Figure 3e). We confirmed that knockout of APP reduces Aβ secretion and that both wild type APP (APP-770) and SPA4CT-T43P are able to rescue Aβ secretion to near physiologic levels (Figure 3f). *In-vivo*, the loss of brain metastasis observed upon APP knockout was rescued by introduction of either APP-770 or SPA4CT-T43P (Figure 3g,h), demonstrating that Aβ is the form of APP required for melanoma brain metastasis.

### Melanoma-secreted Amyloid Beta is required for late growth and survival in the brain parenchyma

To investigate how Aβ functions in melanoma brain metastasis, we began by identifying which step of the brain metastatic process Aβ is required for. First, we established the timeline of brain metastasis in 12-273 BM STC by *ex-vivo* immunofluorescence analysis of brain slices (Figure 4a upper panels, Supplementary Videos 2-6. At day 1 post intracardiac injection, melanoma cells are arrested in the brain microvasculature. By day 3, melanoma cells have extravasated into the parenchyma and remain adhered to the surface of blood vessels with a rounded morphology. From days 3 to 7, cells begin to divide and spread along the vasculature in an elongated morphology, a process known as vascular co-option^8^. By day 7, approximately two thirds of the melanoma cells that had initially reached the brain at day 1 have died. This occurs when melanoma cells either fail to extravasate (Figure 4c) or undergo apoptosis upon entering the brain parenchyma (Figure 4d). From days 7-14, surviving melanoma cells begin to proliferate in a rounded morphology independent of the vasculature to form micro-metastases. After day 14, cells rapidly divide, forming small macro-metastases visible to the naked eye by day 21.

**Figure 4:**
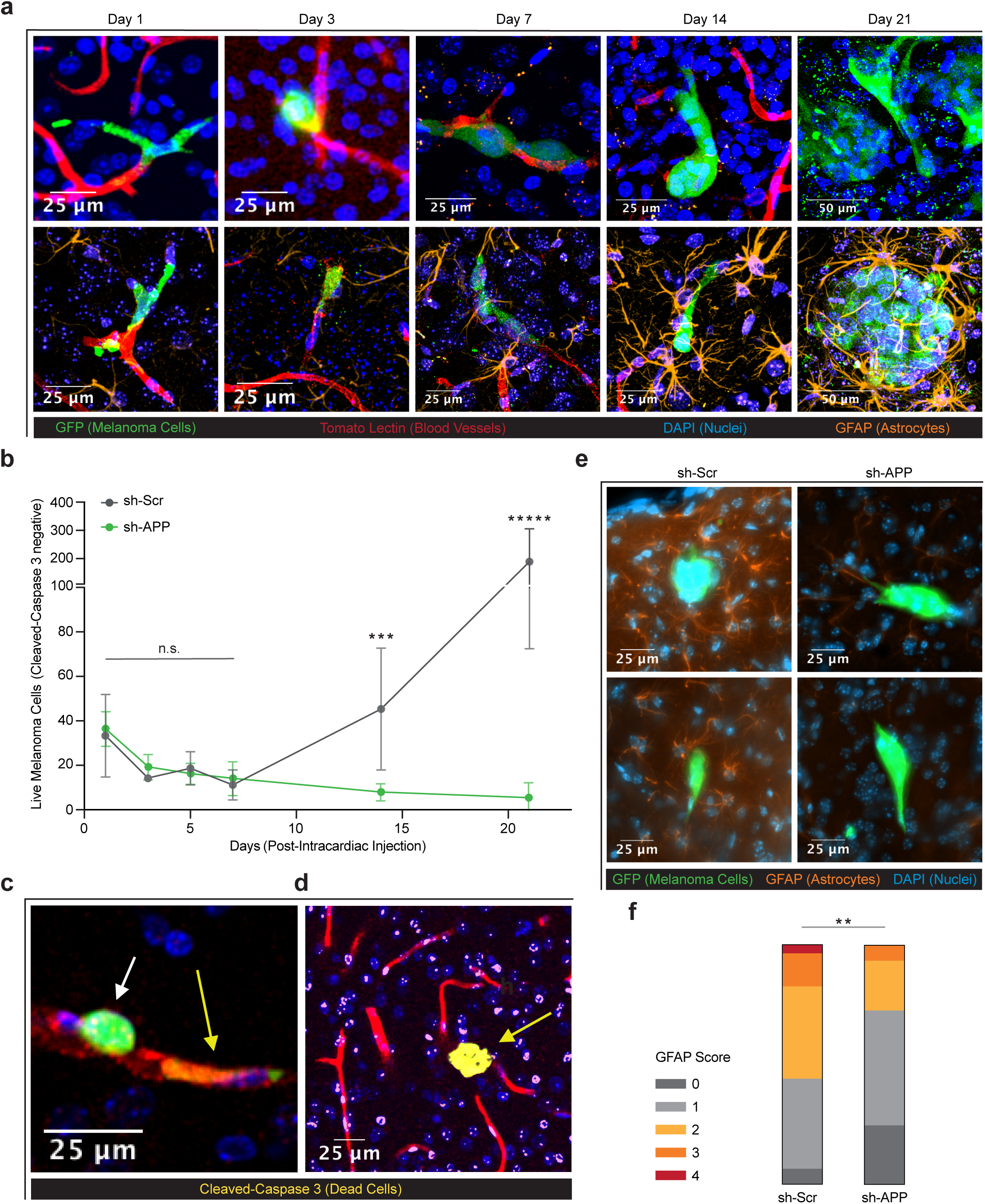
Melanoma-secreted Aβ is Required for Late Growth and Survival in the Brain Parenchyma. **a**, Representative images of brain slice immunofluorescence of 12-273 BM cells at days 1, 3, 7, 14 and 21. Fluorescent markers: green = anti-GFP (melanoma cells), red = tomato lectin (blood vessels), blue = DAPI (nuclei), orange = anti-GFAP (astrocytes). (Images are from 3 mouse brains per group per time point per experiment, 2 independent experiments). **b**, Quantification of live 12-273 BM cells in the brain parenchyma over time after intracardiac injection. Day 14 sh-Scr vs. sh-APP (*** p < .0005), day 21 sh-Scr vs. sh-APP (***** p<.000005). (n= 2 independent experiments, 6 mice per group in total). **c,d**, Images of brain slice immunofluorescence showing live (white arrow) and dead (yellow arrows) cells. Fluorescent marker: yellow = anti-Cleaved-Caspase 3 (dead cells). **e**, Representative images of brain slice immunofluorescence at day 10 post intracardiac injection. **f**, Quantification of qualitative scoring of astrocyte reactivity to live melanoma cells at day 10 post-intracardiac injection. sh-Scr vs sh-APP (chi-square 0-1 = negative, 2-4 = positive ** p<0.005) (n= 2 independent experiments, 4 mouse brains per group, representative experiment shown)

When comparing melanoma cells with and without Aβ, we did not observe differences in the number of live cells from day 1 through day 7 post intracardiac injection (Figure 4b), indicating that Aβ is not required for vascular arrest, extravasation, or early survival of melanoma cells in the brain parenchyma. Melanoma cells lacking Aβ, however, were unable to successfully proliferate to form micro- and macro-metastases by days 14 and 21 respectively (Figure 4b), and instead underwent apoptosis (Extended Data Figure 4a). Therefore, Aβ is required for melanoma cells to progress from vascular co-option to successful metastatic colonization of the brain parenchyma.

### Melanoma-secreted Aβ stimulates local reactive astrocytosis

Given that reactive astrocytes are important regulators of brain metastasis ^10^ and that amyloid beta can influence astrocyte physiology ^26, 27^, we hypothesize that Aβ secreted by melanoma cells triggers a reactive phenotype in surrounding astrocytes that supports melanoma growth in the brain. GFAP staining revealed an increase in the presence of reactive astrocytes surrounding melanoma cells over time (Figure 4a, lower panels), with significant physical contacts between melanoma cells and astrocytes developing from days 7 to 14. By day 21, reactive astrocytes form a glial scar-like structure that envelops the growing brain metastases (Figure 4a, bottom right; Supplementary Video 7). Notably, the time period during which melanoma cells lacking Aβ fail to survive overlaps with the time in which astrocytes begin to extensively interact with melanoma cells. We analyzed astrocytes surrounding live melanoma cell clusters with and without Aβ in the brain at day 10 post-injection by GFAP staining. Clusters of cells lacking Aβ display significantly decreased local reactive astrocytosis than control cells (Figure 4e,f; Extended data figure 4b), indicating that melanoma-secreted Aβ stimulates local reactive astrocytosis.

We further interrogated how melanoma-derived Aβ affects the phenotype of astrocytes by exposing primary rat astrocytes to melanoma-conditioned media. Primary rat astrocytes were isolated and maintained at rest *in-vitro* in serum-free media, as previously reported^28^. We exposed astrocytes to melanoma conditioned media (CM) lacking Aβ, either by genetic silencing of APP in melanoma cells (sh-APP IgG) or by immunodepletion of Aβ from control melanoma CM (sh-Scr Anti-Aβ), and compared them to astrocytes exposed to control melanoma CM (sh-Scr IgG) (Figure 5a). Astrocytes exposed to media with Aβ displayed more elongated branches, a phenotype related to astrocyte reactivity^29^, as compared to those exposed to media without Aβ (Figure 5b,c).

**Figure 5:**
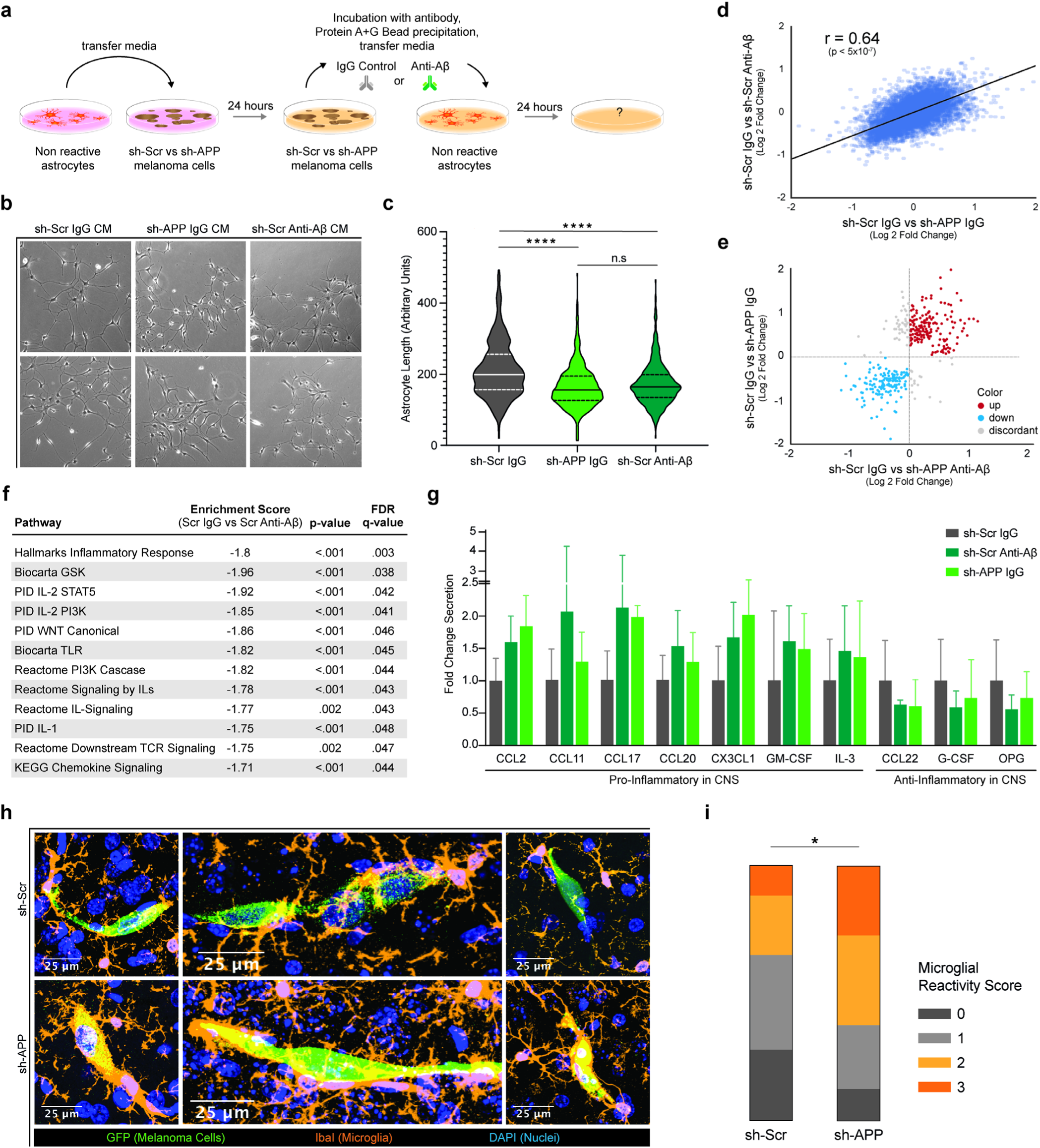
Melanoma-secreted Aβ Induces an Anti-Inflammatory Response in Astrocytes and Inhibits Microglial Activation. a, Diagram of experimental method used to expose astrocytes to melanoma-conditioned media with and without Aβ. b, Representative images of astrocytes after 24-hour exposure to melanoma-conditioned media (CM). c, Quantification of astrocyte length. Sh-Scr IgG CM vs sh-APP IgG CM (**** p<0.00005), sh-Scr IgG CM vs sh-Scr Anti-Aβ (**** p<0.00005). (n = two independent experiments, 4 biological replicates per group, representative experiment shown). d,e Gene expression changes induced in astrocytes upon removal of Aβ from melanoma conditioned media by genetic silencing of APP in melanoma cells (x axis – sh-Scr IgG CM vs sh-APP IgG CM) compared to changes induced by direct immunoprecipitation of Aβ from conditioned media (y axis – sh-Scr IgG CM vs sh-Scr Anti-Aβ CM). f, Enriched pathways identified from GSEA analysis of global gene expression changes in astrocytes exposed to sh-Scr IgG CM vs sh-Scr Anti-Aβ CM. (n = 2 biological replicates per group) g, Quantification of cytokine secretion by cytokine array in astrocytes exposed to melanoma conditioned media (two independent experiments, 4 biological replicates per group, representative experiment shown). h, Representative images of microglia surrounding 12-273 melanoma cells at day 10 post intracardiac injection. Fluorescent markers: green = anti-GFP (melanoma cells), orange= anti-IbaI (microglia), blue = DAPI (nuclei). i, Quantification of qualitative scoring of microglial reactivity to melanoma cells at day 10 post-intracardiac injection. sh-Scr vs sh-APP (chi-square 0-1 = negative, 2-3 = positive * p<0.05). (n = 4 mouse brains per group)

### Melanoma-secreted Aβ induces an anti-inflammatory response in astrocytes

When comparing global transcriptomic changes in astrocytes exposed to melanoma CM with and without Aβ, we found a high degree of correlation (r=0.64) between changes induced by APP silencing in melanoma cells and those induced by Aβ immunodepletion from CM of control melanoma cells (Figure 5d,e). This demonstrates that melanoma-secreted Aβ accounts for the majority of APP-mediated transcriptomic changes that melanoma cells induce in astrocytes, and establishes secreted Aβ as a direct mediator of crosstalk between melanoma cells and astrocytes.

To further characterize how melanoma-derived Aβ influences astrocytes, we performed gene set enrichment analysis of differentially expressed transcripts in astrocytes exposed to melanoma CM with and without Aβ. Intriguingly, results showed a significantly decreased enrichment score in multiple pathways related to inflammatory signaling in astrocytes exposed to media with Aβ as compared to those exposed to media without Aβ (Figure 5f).

Therefore, we hypothesized that a key function of melanoma derived-Aβ could be to stimulate astrocytes to an anti-inflammatory phenotype that helps prevent immunemediated clearance of melanoma cells in the brain parenchyma. Using a cytokine array, the levels of astrocyte-derived secreted factors were quantified in supernatants from astrocytes exposed to melanoma CM in the presence or absence of Aβ. This analysis revealed that Aβ both stimulates astrocyte secretion of anti-inflammatory cytokines and suppresses astrocyte secretion of cytokines with known pro-inflammatory activity in the central nervous system (Figure 5g) ^30–39^, thus demonstrating that melanoma-secreted Aβ stimulates astrocytes to an anti-inflammatory phenotype.

### Aβ suppresses microglia activation and phagocytic clearance of melanoma cells

Several of the identified astrocyte-secreted cytokines have documented roles in microglial chemotaxis ^40^, activation ^31^, and M1 polarization ^41^. We therefore sought to examine the effect of melanoma-secreted Aβ on the recruitment of microglia to the metastatic site. Resident microglia surrounding melanoma cells with and without Aβ were visualized by IbaI immunofluorescent staining in brain slices. Microglia surrounding melanoma cells lacking Aβ exhibited a more ameboid morphology (Figure 5h), which signifies increased microglial activation ^42^. Furthermore, we observed a significantly increase in microglial phagocytosis of melanoma cells lacking Aβ (Figure 5i, Extended data Figure 4c, and Supplementary Video 8), demonstrating that Aβ secretion protects melanoma cells from phagocytic clearance by microglia.

### Aβ is a Promising Therapeutic Target for Treatment of Brain Metastasis

To assess if targeting Aβ could be a promising therapeutic strategy for treatment of melanoma brain metastasis, we investigated whether Aβ is required for growth and survival of established brain metastases. Using a doxycycline inducible shRNA system, we depleted APP in growing melanoma brain macro-metastases (Extended data Figure 5a; Figure 6a). Loss of ability to produce Aβ in pre-existing brain metastases resulted in decreased brain metastatic burden (Figure 6b-d). Many therapeutic approaches efficiently targeting amyloid beta have been developed and tested for the treatment of Alzheimer’s disease, including anti-Aβ antibodies ^43^ and *β*-secretase (BACE) inhibitors ^44^. We tested LY2886721, a BACE inhibitor that blocks Aβ production ^45^ by inhibiting the rate limiting step in its generation (Figure 6e). Treatment of mice with LY2886721 at a dose of 75 mg/kg/day in food resulted in a 75% reduction in plasma Aβ levels (Extended data Figure 5b). Pharmacological inhibition of Aβ production decreased brain metastatic burden in both a patient-derived short-term culture (12-273BM; Figure 6f-h) and an established melanoma cell line (131/4-5B1; Figure 6i-k). Our data support Aβ-targeting as a promising therapeutic approach against melanoma brain metastasis.

**Figure 6:**
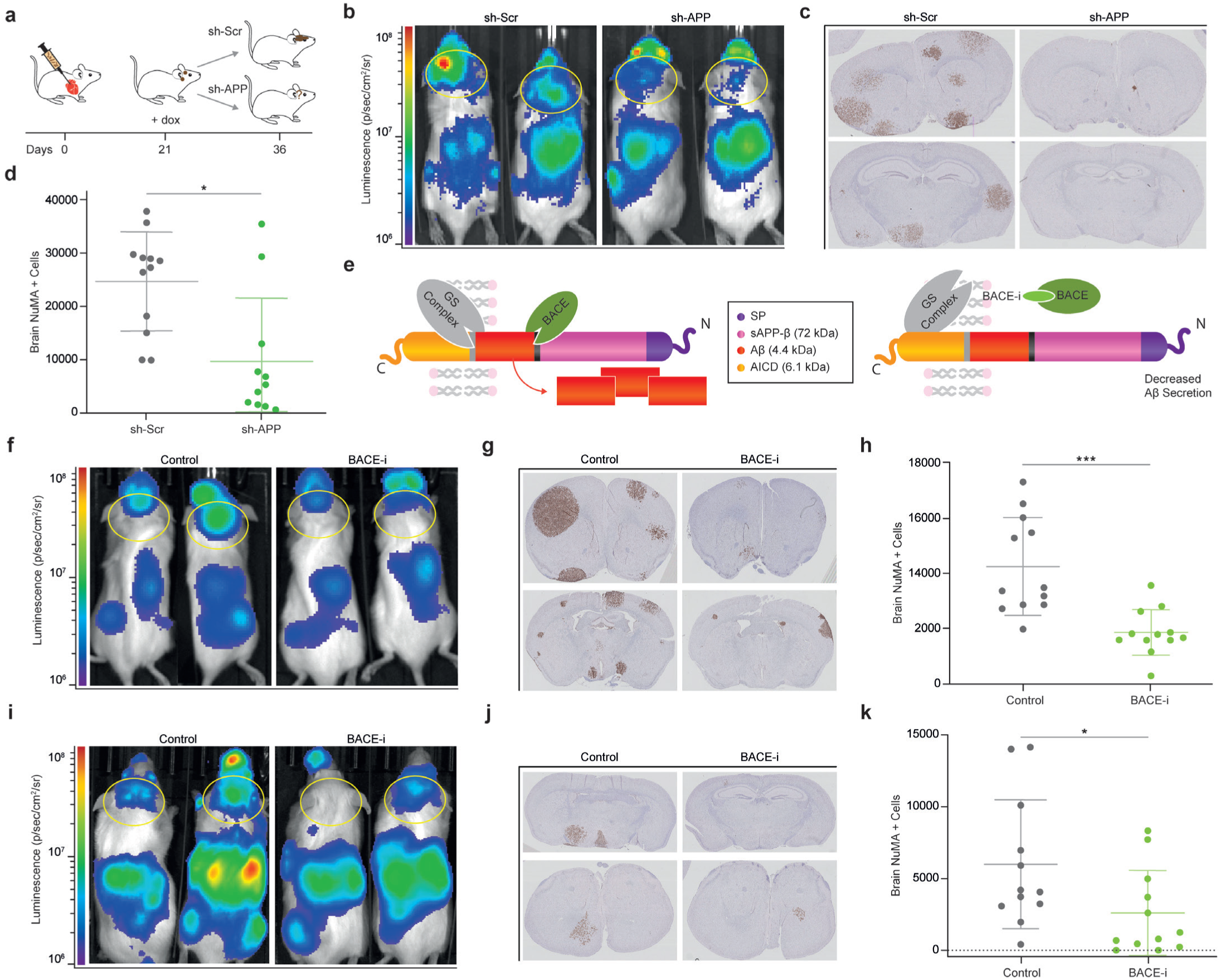
Aβ is a Promising Therapeutic Target for Treatment of Brain Metastasis. **a**, Diagram of therapeutic simulation experiment inducing silencing of APP in established brain metastases. **b**, Representative IVIS images at day 37 post-intracardiac injection. **c**, Representative images of FFPE brain slides with labeling of metastatic cells by anti-NuMA immunohistochemistry. **d**, Quantification of NuMA+ metastatic cells in mouse brains. sh-Scr vs sh-APP (* p<0.05). (n = 10-12 mice per group) **e**, Diagram of APP cleavage and beta-secretase inhibition of Aβ production. **f,i** Representative IVIS images at Day 28 post intracardiac injection with 12-273 BM STC (f) and 5B1 melanoma cell line (**i**). **g,j**, Representative images of FFPE brain slides with labeling of metastatic cells by anti-NuMA immunohistochemistry (**g** – 12-273 BM, **j** – 5B1). **h,k** Quantification of NuMA+ metastatic cells in mouse brains. 12-273 BM Control vs BACE-i (**h** *** p<0.0005), 5B1 Control vs BACE-i (**k** * p<0.05) (n = 10-12 mice per group)

## Discussion

Identifying and characterizing mechanisms that mediate survival of cancer cells in the brain parenchyma has been historically challenging. The majority of studies aiming to identify mechanisms of brain metastasis have utilized cell lines originally derived from extracranial metastases and examined transcriptomic changes after serial transplantation in mice to increase brain tropism ^12, 46–48^. While these models have been an invaluable tool, the process by which they are generated varies greatly from the brain metastatic process occurring in patients ^49^. In contrast, STC pairs exhibiting differential brain tropism (Figure 1b-d and Extended data fig. 1a,b) are derived directly from naturally arising metastases in patients ^50^. Recently, some studies have directly profiled gene expression of surgically resected patient tumors^19, 51^. While this method has several advantages, such as strong clinical relevance and ability to capture effects mediated by the brain microenvironment, the limited amount of material available per sample can restrict the type of analyses that are technically feasible. Furthermore, direct use of patient biopsies makes it difficult to establish whether genes identified as upregulated in CNS metastases represent contamination from brain tissue or neural mimicry by cancer cells. Proteomic screening of short term cultures – which no longer contain normal brain tissue – circumvents this issue and allowed us to identify an association between brain metastasis and proteins implicated in Alzheimer’s disease.

Several studies have demonstrated that reactive astrocytes support the formation of brain metastasis ^10, 11^, provided early antagonistic interactions can be overcome ^12^. Here, we show that Aβ secreted by melanoma cells stimulates astrocytes to exhibit a pro-metastatic, anti-inflammatory phenotype. It was demonstrated that cancer cells in the brain induce a subpopulation of pro-metastatic Stat3 positive reactive astrocytes in their microenvironment by an unknown mechanism ^52^. In addition, Stat3^+^ reactive astrocytes contribute to the pathology of acute brain injury and several neurodegenerative diseases, including Alzheimer’s disease. Further investigation is warranted to address whether melanoma-secreted Aβ specifically induces Stat3 activation in astrocytes.

Astrocytes have been shown to regulate microglial activity in response to inflammatory insults ^53^. A similar role for astrocytes has been theorized in the brain metastatic microenvironment but remains largely unexplored ^54^. Here, we establish that melanoma-secreted Aβ directly suppresses astrocyte secretion of several cytokines that recruit and activate microglia, such as CCL2, a potent chemoattractant that polarizes microglia to a pro-inflammatory, anti-tumorigenic M1 phenotype ^41^. Furthermore, we demonstrate that melanoma-secreted Aβ inhibits microglial phagocytosis of melanoma cells. Taken together, these results suggest that astrocytes regulate microglial activity in the brain metastatic microenvironment. It is also possible that melanoma-secreted Aβ impacts microglial response to melanoma cells directly. A recent study showed that acute exposure to Aβ initially stimulated microglial phagocytosis of microparticles, but re-exposure of the same microglia to Aβ days later instead inhibited phagocytosis^55^. Given the high Aβ concentrations used and lack of a continuous exposure of microglia to Aβ, it is difficult to interpret that study’s findings in the context of the brain metastatic microenvironment. Additional studies are needed to clarify whether melanoma-secreted Aβ directly impacts microglial function.

The role of Aβ, both in Alzheimer’s disease and in normal physiology, remains controversial. Most studies investigating the function of Aβ have been performed in the context of Alzheimer’s pathology using transgenic mice that mimic the human disease ^56^. It is well established that insoluble aggregates of Aβ contribute to pathological astrocyte- and microglia-mediated neuroinflammation in Alzheimer’s disease ^57^. Furthermore, Aβ oligomers have been shown to act as anti-microbial peptides and protect against CNS infection ^58, 59^. Intriguingly, we demonstrate an anti-inflammatory function of soluble Aβ in the context of brain metastasis. The majority of Aβ produced by melanoma cells is Aβ-40 (Figure 3 c,f, data not shown), the less aggregative form, and none of our models of brain metastasis gives rise to Aβ plaques (data not shown). Instead, we hypothesize that melanoma-derived Aβ acts in the form of soluble monomers or oligomers to suppress astrocyte-driven inflammation. Indeed, both monomers and oligomers of Aβ have been shown to affect phenotypic changes in astrocytes ^26, 27^.. Whether soluble Aβ also acts as an anti-inflammatory mediator in the brain during normal physiology and other pathophysiologic contexts is an important question that requires further investigation. If present, a similar function of soluble Aβ could profoundly impact our understanding of Alzheimer’s development and shed light on the reported lack of efficacy of anti-Aβ agents against advanced Alzheimer’s disease.

Our studies show Aβ is a highly promising therapeutic target for melanoma brain metastasis. The brain represents an immune privileged environment and is often a site of treatment resistance or relapse in patients ^4^. Although checkpoint blockade immunotherapy, the standard-of-care for metastatic melanoma, can be efficacious in brain metastasis, the rates of on-treatment progression are higher for intracranial than for extracranial metastases^5^. Targeting Aβ, which suppresses neuroinflammation, in combination with immune checkpoint inhibitors could result in a more robust anti-tumor immune response and improve patient outcomes. Several therapeutic agents targeting Aβ have been developed and extensively tested in clinical trials for treatment of Alzheimer’s disease and could be repurposed for treatment of melanoma brain metastasis. One possibility includes BACE small molecule inhibitors, exemplified by LY2886721, which provided proof-of-principle efficacy in our preclinical models. Another attractive approach is the use of anti-Aβ antibodies, which have been extensively tested in clinical trials for Alzheimer’s disease. Anti-Aβ antibodies successfully sequestered Aβ in Alzheimer’s patients ^60^ but lacked clinical efficacy ^61^. In Phase III clinical trials, anti-Aβ antibodies were tolerated at high doses for extended periods of time without an increase in rates of adverse effects over placebo ^61^. Given that dose limiting toxicities and adverse effects are major drivers of failure in cancer clinical trials, targeting Aβ in melanoma brain metastasis with anti-Aβ antibodies represents a particularly promising treatment strategy.

In summary, our studies reveal an unexpected role for tumor-secreted Aβ, a polypeptide heavily implicated in Alzheimer’s disease, in the adaptation of melanoma cells to the brain microenvironment and provide proof-of-principle of Aβ targeting as a novel therapeutic avenue against this devastating condition.

---

**Extended Data Figure 1:**
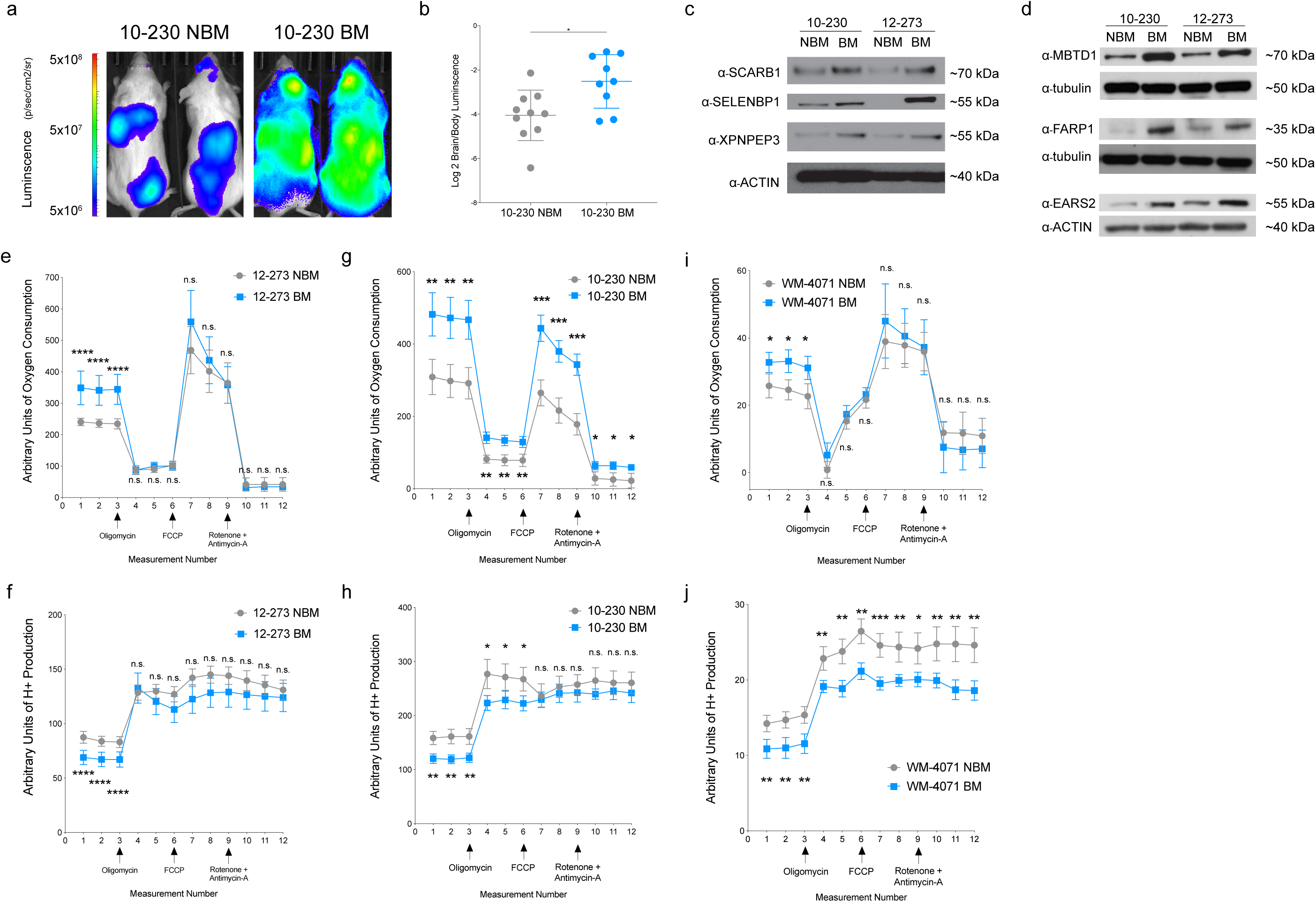
**a**, Representative IVIS images of 10-230 STC pair at 35 days post-intracardiac injection in mice. **b**, Quantified brain/body luminescence ratio on day 35. 10-230 NBM vs BM (* p<0.05). (n = 2 independent experiments, 10 mice per group, representative data shown). **c,d** Western blot analysis of STC pairs for differentially expressed proteins identified in proteomics. **e-j**, Seahorse metabolic analysis of STC pairs. 12-273 pair oxygen consumption (**e**, **** p<0.00005) and H+ production (**f**, **** p<0.00005), 10-230 pair oxygen consumption (**g**, ** p<0.005) and H+ production (**h**, ** p<0.005), WM-4071 pair oxygen consumption (**i**, * p<0.05) and H+ production (**j**, ** p<.005). (10-230 pair, 12-273 pair, n = 3 independent experiments, 4-6 biological replicates per group, representative experiment shown. WM 4071 pair n= 1 experiment, 4-6 biological replicates per group)

**Extended Data Figure 2:**
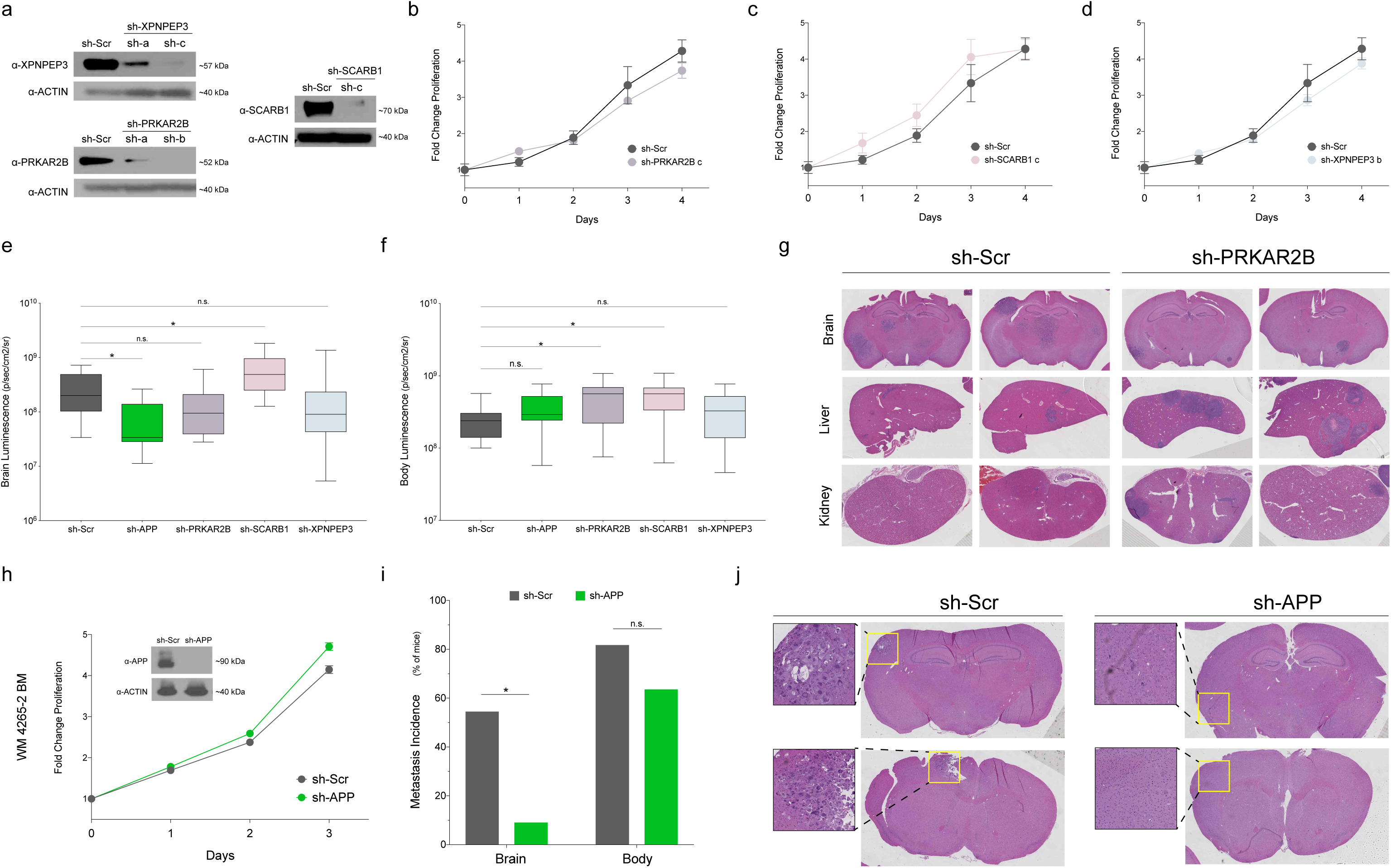
**a**, Western blot of analysis of 12-273 BM cells transduced with the indicated sh-RNA carrying lentivirus. The following sh-RNAs were used in the miniscreen: sh-XPNPEP3-c, sh-PRKARB-c, sh-SCARB1-c. **b-d**, Fold change in-vitro proliferation and western blot analysis of 12-273BM cells with (**b**) sh-Scr vs sh-PRKAR2B, (**c**) sh-Scr vs sh-SCARB1, (**d**) sh-Scr vs sh-XPNPEP3. **e**, Quantified brain luminescence in mice 35 days post-intracardiac injection of 12-273 BM with shRNA-mediated silencing of selected candidates or sh-Scr. sh-Scr vs sh-APP (* p<0.05), sh-Scr vs sh-SCARB1 (* p<0.05). (Box = Interquartile range. Error bars = min to max.) **f**, Quantified body luminescence in mice 35 days post-intracardiac injection of 12-273 BM with shRNA-mediated silencing of selected candidates or sh-Scr. sh-Scr vs sh-PRKAR2B ( * p<0.05), sh-Scr vs sh-SCARB1 (* p<0.05) (Box = Interquartile range. Error bars = min to max.) **g**, Representative H&E-stained FFPE sections of brains, livers, and kidneys from mice injected with 12-273 BM sh-Scr and sh-PRKAR2B. Each column contains images from the same mouse. (n= 1 experiment, 10-12 mice per group). **h**, Fold change in-vitro proliferation and western blot analysis of WM 4265-2 BM cells with sh-Scr vs sh-APP. **i**, Incidence of brain and body metastasis in mice injected with WM 4265-2 BM at 88 days post-intracardiac injection. sh-Scr vs sh-APP brain (chi square, * p<0.05). **j**, H&E stained FFPE sections of brains from mice injected with WM-4265-2 BM sh-Scr and sh-APP. Upper sh-APP image is from the single sh-APP mouse with brain metastasis. (n = 10-12 mice per group). **k**, Video of 3D projections from MRI images of representative of mouse brains. Pink-purple – brain metastasis (12-273 BM). Top row: sh-Scr. Bottom row: sh-APP.

**Extended Data Figure 3:**
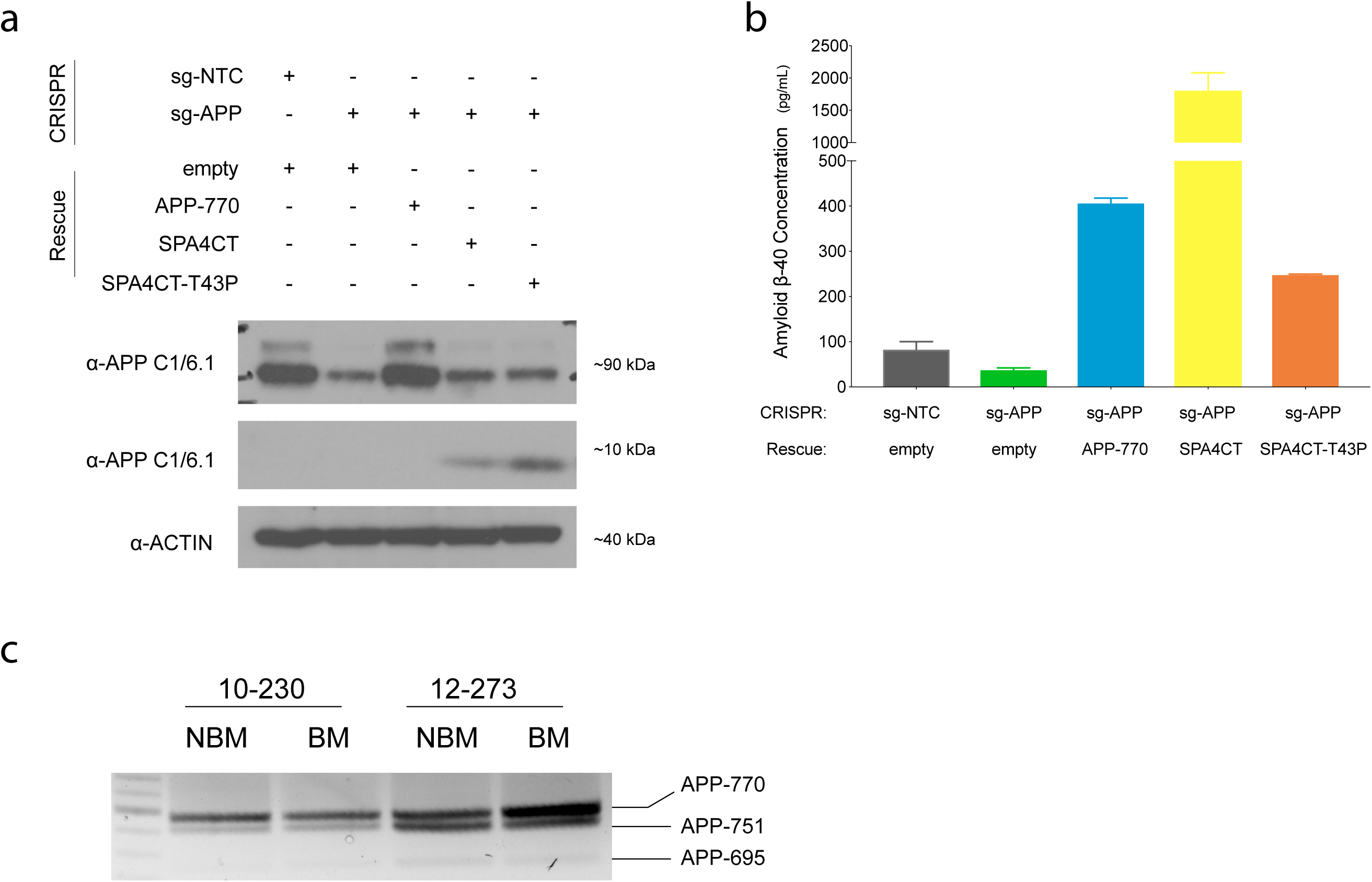
**a**, Western blot analysis of 12-273 BM infected cells for APP and SPA4CT. **b**, Quantification of Aβ secretion by ELISA in 12-273 BM infected cells. **c**, Semi-quantitative RT-PCR of APP isoforms in 10-230 and 12-273 STC pairs.

**Extended Data Figure 4:**
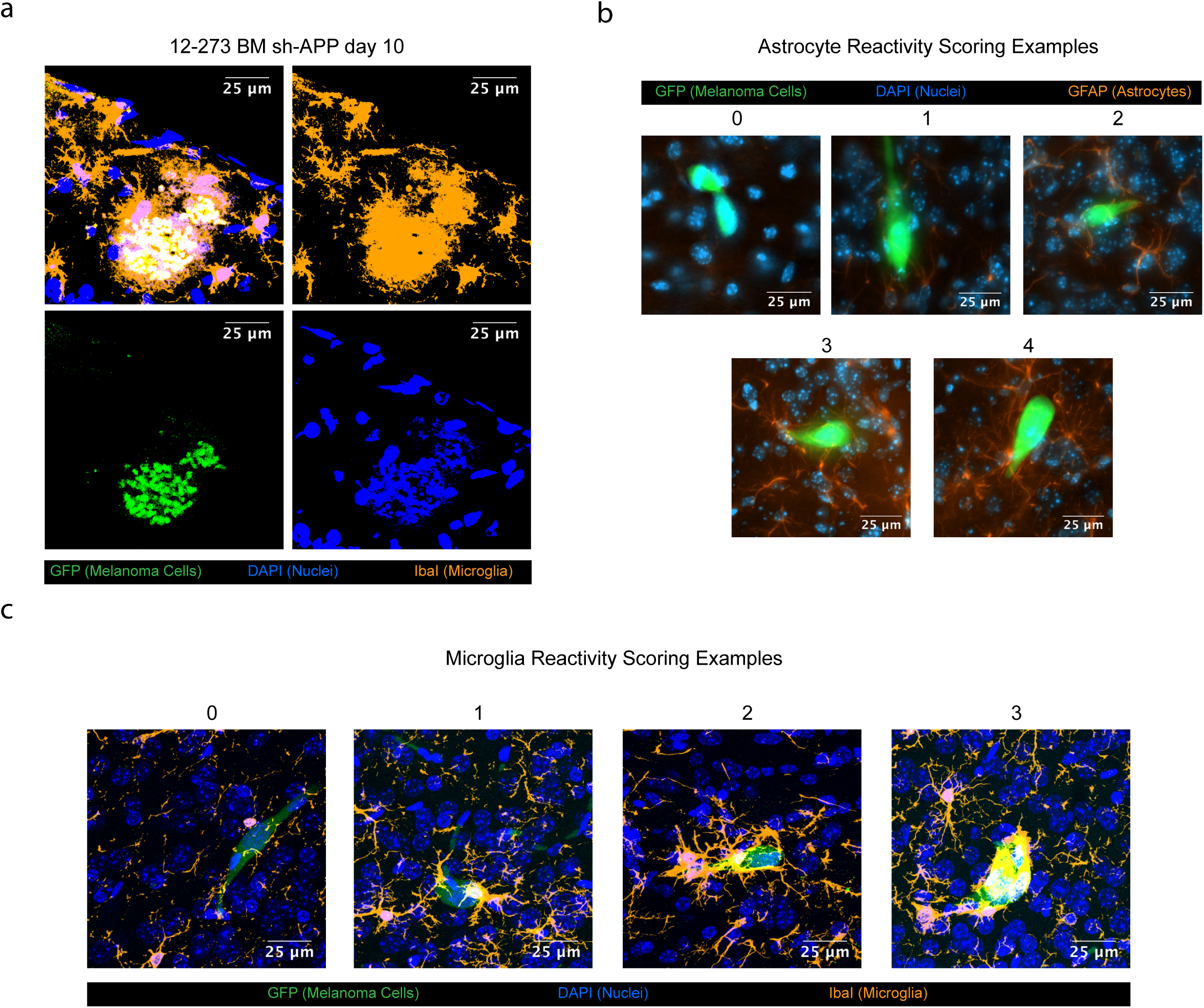
**a**, Image of microglial phagocytosis of apoptotic bodies from brain slices of mice instilled with 12-273 BM sh-APP cells 10 days post-intracardiac injection. Fluorescent markers: green = anti-GFP (melanoma cells), blue = DAPI (nuclei), orange = anti-IbaI (microglia). **b**, Representative images of astrocyte reactivity scores. Fluorescent markers: green = anti-GFP (melanoma cells), blue = DAPI (nuclei), orange = anti-GFAP (astrocytes). **c**, Representative images of microglia reactivity scores. Fluorescent markers: green = anti-GFP (melanoma cells), blue = DAPI (nuclei), orange = anti-IbaI (microglia). **d-i**, 3D projection of confocal images of melanoma cells in brain parenchyma at Day 1 (**d**), Day 3 (**e**), Day 7 (**f**), Day 14 (**g**), and Day 21 (**h-i**). Fluorescent markers: green = anti-GFP (melanoma cells), blue = DAPI (nuclei), orange = anti-GFAP (astrocytes).

**Extended Data Figure 5.**
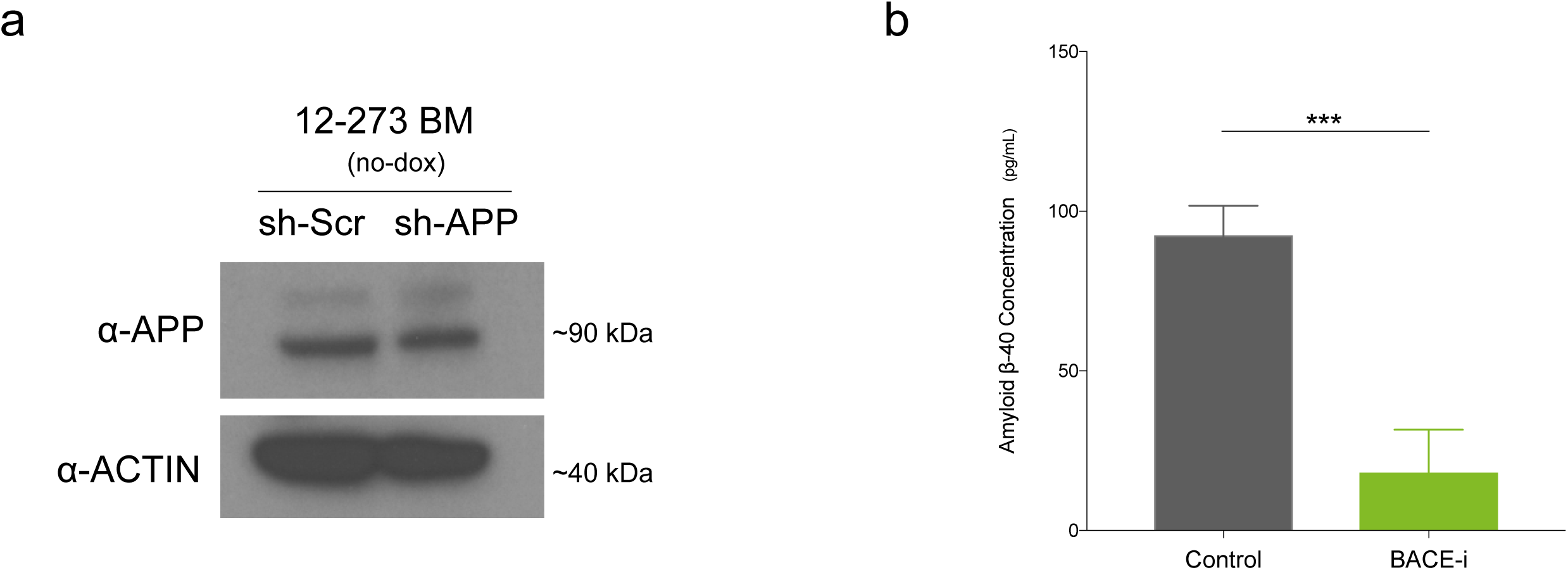
**a**, Western blot analysis of 12-273 BM sh-Scr and sh-APP cells cultured without doxycycline on day of intracardiac injection. **b**, Quantification of plasma Aβ levels in Control vs BACE-i treated mice at 28 days post-intracardiac injection with 12-273 BM cells.

## Supplementary Videos

**Supplementary video 1** – Representative 3D renderings from MRI images of brains from mice at 35 days after intracardiac injection with 12-273 BM. Pink/purple = metastasis. Top row: sh-Scr. Bottom row: sh-APP.

**Supplementary Videos 2-6 –** 3D renderings from confocal images of 12-273 BM sh-Scr cells in mouse brain parenchyma at the following times after intracardiac injection: day 1 (Video 2), day 3 (Video 3), day 7 (Video 4), day 14 (Video 5), and day 21 (Video 6). Fluorescent markers: green =anti-GFP (melanoma cells), red = tomato lectin (blood vessels), blue = DAPI (nuclei).

**Supplementary Video 7 –** 3D rendering from confocal image of 12-273 BM sh-Scr cells in mouse brain parenchyma at 21 days after intracardiac injection. Fluorescent markers: green = anti-GFP (melanoma cells), red = tomato lectin (blood vessels), blue = DAPI (nuclei), orange = anti-GFAP (astrocytes).

**Supplementary Video 8 –** 3D rendering from confocal image of 12-273 BM sh-APP cells at 10 days after intracardiac injection. Video depicts melanoma cells that have been completely phagocytized by microglia. Fluorescent markers: green = anti-GFP (melanoma cells), orange= anti-IbaI (microglia), blue = DAPI (nuclei)

## Methods

### Cell culture

#### Melanoma short-term cultures and cell lines

Low passage melanoma short-term cultures (STCs), derived in the Osman laboratory as described ^50^. were grown in DMEM with 10% fetal bovine serum (FBS), 1 mM Sodium Pyruvate, 4 mM L-Glutamine, 25 mM D-Glucose, 1% Nonessential Amino Acids (NEAA), 100 units/mL penicillin, and 100 ug/mL streptomycin. Additional STCs, a kind gift of the Herlyn lab at the Wistar Institute, were grown in Tu2% media (80% MCDB153, 20% Leibovitz’s L-15, 2% FBS, 5ug/ml Insulin (Bovine), 1.68 mM calcium chloride, 100 units/mL penicillin, 100 ug/mL streptomycin). STCs were kept below lifetime passage number of 40 for all experiments. For details of STCs utilized, see Extended Data Table 1. HEK293T cells (for lentivirus production) and 131/4-5B1 (hereafter 5B1, ^46^) melanoma cells were grown in DMEM with 10% fetal bovine serum (FBS), 1 mM Sodium Pyruvate, 4 mM L-Glutamine, 25 mM D-Glucose, 100 units/mL penicillin, and 100 ug/mL streptomycin. Cell lines were maintained in a 5% CO2 incubator at 37 °C and were routinely tested for Mycoplasma contamination.

#### Astrocytes isolation and culture

Astrocytes were purified by immunopanning and cultured in serum-free conditions as previously described ^62^. Briefly, cortices from 5-6 postnatal day 4-6 Sprague Dawley rat pups (Charles River) were dissected out and meninges and choroid plexus removed. The cortices were minced with a scalpel and digested in Papain for 40 min at 34°C under constant CO_2_/O_2_ gas equilibration. The digested brain pieces were washed with CO_2_/O_2_-equilibrated Ovomucoid inhibitor solution, triturated, and spun down through a cushion gradient containing low and high Ovomucoid inhibitor layers. The resulting cell pellet was passed through a 20 µm nylon mesh to create a single cell suspension. The cells were then incubated in a 34°C water bath for 30-45 mins to allow cell-specific antigens to return to the cell surface. Negative selection was performed using Goat anti-mouse IgG + IgM (H + L), *Griffonia (Bandeiraea) simplicifolia* lectin 1 (BSL-1), Rat anti-mouse CD45, and O4 hybridoma supernatant mouse IgM ^63, 64^, followed by positive selection for astrocytes using mouse anti-human integrin β5 (ITGB5). Purified astrocytes were detached from the panning plate with trypsin at 37°C for 3-4 min, neutralized by 30% fetal calf serum, counted, pelleted, and resuspended in 0.02% BSA in DPBS. All isolation and immunopanning steps occurred at room temperature, except for the heated digestion, incubation, and trypsinization steps. Cells were plated at 50,000 or 70,000 cells per well in 6 well plates containing 2 mL/well of serum-free Astrocyte Growth Medium (50% Neurobasal Medium, 50% Dulbecco’s Modified Eagle Medium (DMEM), 100 U/mL Penicillin, 100 µg/mL Streptomycin, 1 mM Sodium pyruvate, 292 µg/mL L-Glutamine, 5 µg/mL *N*-Acetyl-L-cysteine (NAC), 100 µg/mL BSA, 100 µg/ml Transferrin, 16 µg/mL Putrescine dihydrochloride, 60 ng/mL (0.2 µM) Progesterone, and 40 ng/mL Sodium selenite. Immediately before plating, the astrocyte trophic factor Heparin-binding EGF-like growth factor (HBEGF) was added (5 ng/mL) and media equilibrated to 37°C in a 10% CO_2_ incubator). Cells were incubated at 37°C in 10% CO_2_. Media was changed (50% well volume) once per week.

#### Exposure of Astrocytes to Melanoma Conditioned Media

Astrocytes were plated at 5×10^4^ astrocytes/well in 6-well plates. 12-273 BM melanoma cells were plated at 2.5×10^5^ cells/well in 6-well plates. Astrocyte-conditioned media (ACM) was obtained by culture of astrocytes in astrocyte media for 1 week. 1.5mL of ACM were then transferred and incubated with 12-273 BM cells in 6-well plates for 24 hours. Media was then removed and incubated with either 4.5 ug of mouse IgG or 2.25 ug of anti-amyloid beta antibody N25 and 2.25ug of anti-APP 6E10 antibody for 1 hr at 4 degrees and 1 hr at room temperature with rotation mixing. 75ul of protein A/G beads (Pierce 88803) were washed twice with PBS, added to 1.5mL of conditioned media, and incubated for 1 hour at room temperature with rotation mixing. Beads were precipitated using a magnetic tube holder and 1.2 mL of conditioned media was transferred per well to astrocytes in a 6-well plate. Astrocytes were incubated for 24 hours.

#### Astrocyte Length Quantification

After 24-hour incubation in melanoma conditioned media, pictures of astrocytes were randomly captured on light microscopy at 10x. Average astrocyte length was quantified by measurement of longest dimension of each astrocyte using ImageJ.

#### Astrocyte RNA-Seq and Analysis

After 24-hour incubation in melanoma conditioned media, RNA was harvested from astrocytes using the RNeasy Mini Kit (Qiagen 74104). RNA-Seq library preps were made using the Illumina TruSeq® Stranded mRNA LT kit (Cat #RS-RS-122-2101 or RS-122-2102), on a Beckman Biomek FX instrument, using 100 ng of total RNA as input, amplified by 12 cycles of PCR, and run on an Illumina 4000 as single read 50. FastQC v0.11.7 (http://www.bioinformatics.babraham.ac.uk/projects/fastqc/) was used to check fastq files for poor sequencing quality; all samples had high quality. Illumina adapter sequences and poor-quality bases were then trimmed using trimmomatic v0.36^65^. Trimmed sequences were mapped to mm10 using STAR v2.6.0a^66^, indexed using samtools v1.9^67^, then quantified for UCSC genes using HTSeq-count v0.11.1^68^. Comparative analysis between conditions was performed using DESeq2 v1.24.0 ^69^ with default parameters.

#### Measurement of Astrocyte-Secreted Cytokines

After 24-hour incubation in melanoma conditioned media, media was removed from astrocytes and levels of cytokines were measured using the Rat XL Cytokine Array Kit (R&D Systems ARY030) using 1 mL of conditioned media per membrane. Using ImageJ, mean integrated density from spots representing different cytokines was quantified. For each detected cytokine, mean densities of sh-Scr anti-AB and sh-APP IgG were normalized to the mean sh-Scr IgG density and plotted as a mean fold change.

### Proteomics Analysis of Short-Term Cultures (STCs)

#### Protein isolation

STCs at ∼80% confluence at ∼24 hours post-media change were scraped from 10 or 15cm plates on ice, washed once with cold PBS, and lysed in cold RIPA buffer supplemented with protease inhibitor, for 15 minutes with vortexing every 5 minutes. Protein concentration was determined using Micro BCA Protein Assay Kit (Thermo Scientific 23235).

150 µg of each protein lysate were proteolytically digested and subjected to quantitative mass spectrometry on an Orbitrap Fusion Lumos mass spectrometer using isobaric tandem mass tags (TMT) similar to previous studies ^70, 71^.

#### Sample preparation for mass spectrometry analysis

150 µg of each protein lysate were prepared using the filter-aided sample preparation (FASP) method ^72^. Briefly, each sample was reduced with DTT (final concentration of 20 mM) at 57°C for 1 hour and loaded onto a MicroCon 30-kDa centrifugal filter unit (Millipore) pre-equilibrated with 200 μl of FASP buffer [8 M urea and 0.1 M tris-HCl (pH 7.8)]. Following three washes with FASP buffer, lysates were alkylated on a filter with 50 mM iodoacetamide for 45 min in the dark. Filter-bound lysates were washed three times each with FASP buffer followed by 100 mM ammonium bicarbonate (pH 7.8). The samples were digested overnight at room temperature with trypsin (Promega) at a 1:100 ratio of enzyme to protein. Peptides were eluted twice with 100 μl of 0.5 M NaCl. The tryptic peptides were subsequently desalted using an UltraMicro Spin Column, C18 (Harvard Apparatus) and concentrated in a SpeedVac concentrator.

#### TMT labeling

The dried peptide mixture was re-suspended in 100 μl of 100 mM Triethylammonium bicarbonate (TEAB) (pH 8.5). Each sample was labeled with TMT reagent according to the manufacturer’s protocol. In brief, each TMT reagent vial (0.8 mg) was dissolved in 41 μL of anhydrous ethanol and was added to each sample. The reaction was allowed to proceed for 60 min at room temperature and then quenched using 8 μL of 5% (w/v) hydroxylamine. The samples were combined at a 1:1 ratio and the pooled sample was subsequently desalted using strong-cation exchange and strong-anion exchange solid-phase extraction columns (Strata, Phenomenex) as described^2^.

#### Global Proteome Analysis

A 500 μg aliquot of the pooled sample was fractionated using basic pH reverse-phase HPLC (as described) ^71^. Briefly, the sample was loaded onto a 4.6 mm × 250 mm Xbridge C18 column (Waters, 3.5 μm bead size) using an Agilent 1260 Infinity Bio-inert HPLC and separated over a 90 min linear gradient from 10 to 50% solvent B at a flow rate of 0.5 ml/min (Buffer A = 10 mM ammonium formate, pH 10.0; Buffer B = 90% ACN, 10 mM ammonium formate, pH 10.0). A total of 120 fractions were collected and non-concatenated fractions combined into 40 final fractions. The final fractions were concentrated in the SpeedVac and stored at −80 °C until further analysis.

#### LC-MS/MS analysis

An aliquot of each final fraction was loaded onto a trap column (Acclaim® PepMap 100 pre-column, 75 μm × 2 cm, C18, 3 μm, 100 Å, Thermo Scientific) connected to an analytical column (EASY-Spray column, 50 m × 75 μm ID, PepMap RSLC C18, 2 μm, 100 Å, Thermo Scientific) using the autosampler of an Easy nLC 1000 (Thermo Scientific) with solvent A consisting of 2% acetonitrile in 0.5% acetic acid and solvent B consisting of 80% acetonitrile in 0.5% acetic acid. The peptide mixture was gradient eluted into the Orbitrap Lumos Fusion mass spectrometer (Thermo Scientific) using the following gradient: 5%-23% solvent B in 100 min, 23% - 34% solvent B in 20 min, 34%-56% solvent B in 10 min, followed by 56%-100% solvent B in 20 min. Full scans were acquired with a resolution of 60,000 (@ *m*/*z* 200), a target value of 4e5 and a maximum ion time of 50 ms. After each full scan the most intense ions above 5E4 were selected for fragmentation with HCD using the “Top Speed” algorithm. The MS/MS were acquired in the Orbitrap with a resolution of 60,000 (@ *m*/*z* 200), isolation window of 1.5 *m*/*z*, target value of 1e5, maximum ion time of 60 ms, normalized collision energy (NCE) of 35, and dynamic exclusion of 30 s.

#### Data analysis

The MS/MS spectra were searched against the UniProt human reference proteome with the Andromeda ^73^ search engine integrated into the MaxQuant ^74^ environment (version 1.5.2.8) using the following settings: oxidized methionine (M), TMT-labeled N-term and lysine, acetylation (protein N-term) and deamidation (asparagine and glutamine) were selected as variable modifications, and carbamidomethyl (C) as fixed modifications; precursor mass tolerance was set to 10 ppm; fragment mass tolerance was set to 0.01 Th. The identifications were filtered using a false-discovery rate (FDR) of 0.01 using a target-decoy approach at the protein and peptide level. Only unique peptides were used for quantification and only proteins with at least two unique peptides were reported. Data analysis was performed using Perseus. Protein levels were median centered and log2 normalized. To identify proteins differentially expressed proteins between the BM and NBM cohorts, a Welch’s t-test was performed between unpaired 14 BM and 11 NBM unpaired samples and a paired t-test was performed on the 3 sample pairs. Top differentially expressed genes (defined here as p-value < 0.05) were assessed at the pathway level using DAVID GO-term enrichment with an FDR < 0.001 (ref).

### Animal Studies

All experiments were conducted following protocols approved by the NYU Institutional Animal Care Use Committee (IACUC) (protocol number s16-00051). NOD/SCID/IL2yR^-/-^ male mice (Jackson Labs, Cat# 05557) at 6-8 weeks were used for in-vivo studies.

#### Long-term brain metastasis assays

1 × 10^5^ 12-273 NBM, 1 × 10^5^ 12-273 BM, 1 × 10^5^ 10-230 NBM, 1 × 10^5^ 10-230 BM, 2 × 10^5^ 5B1 cells, or 1.5 × 10^5^ WM-4265 BM cells suspended in 100 ul of PBS were injected with ultrasound guidance (Visualsonics Vevo 770 Ultrasound Imaging System) into the left ventricle of mice anesthetized with isoflurane. Mice were monitored weekly for metastatic progression by in-vivo Bioluminescent imaging (BLI). Upon substantial weight loss and/or signs of distress (neurological signs, abnormal locomotion) in any experimental mice, experimental endpoint was established. At experimental endpoint, all mice in all experimental groups were euthanized by perfusion.

For all long-term brain metastasis assays, 12 mice per group underwent intracardiac injection with cancer cells. Experimental group sizes vary from 10 to 12 mice due to infrequent instances of unsuccessful intracardiac injection.

#### In-vivo BLI

10 minutes prior to imaging, luciferin substrate (150mg/kg) was administered to mice by intraperitoneal injection. Mice were anesthetized with isoflurane and imaged by IVIS Illumina instrument (PerkinElmer) for an automatically-determined duration (1-120 sec). Signal was quantified by measurement of total luminescent flux (p/sec/cm^2^/sr) in drawn brain and body regions of interest.

#### Perfusion

Mice were anesthetized with a ketamine (100mg/kg) and xylazine (10 mg/kg) cocktail by intraperitoneal injection. The heart was exposed by gross dissection and an incision was made in the right atrium. Subsequently, 10 mL of PBS followed by 10 mL of 4% PFA was injected into the left ventricle.

#### Short-term brain metastasis assays and brain slice immunofluorescence

1×10^5^ (for live melanoma cell quantification) or 5 × 10^5^ (for astrocyte and microglial scoring) 12-273 BM cells were introduced by intracardiac injection. Mice were euthanized by perfusion at specified time points post-intracardiac injection. Where relevant, 100 ug of Dylight 647 labeled Lycopersicon Esculentum (Tomato Lectin) were injected into the left ventricle 3 min prior to perfusion. For these assays, 3-6 mice with successful intracardiac injection were used per group per time point.

#### Drug treatment experiments

Protocol for long-term brain metastasis assay (as described above) was performed. LY288671 (75mg/kg/day) was administered to mice in food pellets starting one week prior to intracardiac injection of cancer cells and continuing through the experimental endpoint.

#### Continuous shRNA-mediated gene silencing

Protocol for long-term brain metastasis assay (as described above) was performed. Doxycycline hyclate (200mg/kg/day) was administered to mice in food pellets starting two days prior to intracardiac injection of cancer cells and continuing for the experimental duration.

#### Induction of gene silencing in growing metastases

Protocol for long-term brain metastasis assay (as described above) was performed. Mice were administered a normal diet at the beginning of the experiment. 21 days after intracardiac injection of cancer cells, gene silencing was induced by administration of Doxycycline hyclate (200mg/kg/day) in food pellets.

### Mouse tissue processing, histology and NuMA Immunohistochemistry

Organs harvested from PFA-perfused mice were fixed in 10% formalin for 48 hours. Prior to embedding, brains were sectioned grossly into thirds coronally and livers were sectioned by lobe. Organs were embedded in paraffin and cut into 5uM thick sections.

#### Histological Analysis Comparing Short-Term Culture Pairs

The embedded brain thirds were sectioned coronally through the entire length of tissue at an interval of 50uM. All sections were stained with H&E. Number of unique metastases present per brain was assessed by a pathologist by identifying and tracking metastases through serial sections, ensuring each metastasis was counted only once. Slides were blinded to pathologist (R. R.) prior to analysis.

#### Sectioning and NuMA Immunohistochemistry

For kidney and liver, sections were obtained from one and two representative levels respectively. One section per level was stained with H&E. For brain, sections were obtained from 2 (12-273 BM) or 4 (5B1) evenly space levels of the embedded coronal thirds. In total, this resulted in 6(12-273 BM) or 24 (5B1) serial coronal brain section levels for metastatic quantification. One section per level was stained with H&E. Chromogenic Immunohistochemistry was performed on a Ventana Medical Systems Discovery XT instrument with online deparaffinization and using Ventana’s reagents and detection kits unless otherwise noted. Unconjugated, polyclonal rabbit anti-human Nuclear Mitotic Apparatus Protein (NuMA; Abcam Cat# 97585 Lot# B115626 RRID: AB_1855299) was used for labeling. Sections were deparaffinized in xylene and rehydrated in graded ethanol followed by rinsing in deionized water. Epitope retrieval was performed in a 1200-Watt microwave oven at 100% power in 10 mM sodium citrate buffer, pH 6.0 for 10 minutes. NuMA antibody was diluted 1:7000 in Tris-BSA (25 mM Tris, 15 mM NaCL, 1% BSA, pH 7.2) and incubated for 12 hours. Primary antibody was detected with goat anti-rabbit HRP conjugated multimer, and the complexes were visualized with 3,3 diaminobenzidene and enhanced with copper sulfate. Slides were washed in distilled water, counterstained with hematoxylin, dehydrated and mounted with permanent media. Appropriate positive and negative controls were run in parallel to study sections.

#### NuMA+ Cell Quantification

Scanned images of slides were analyzed using Viziopharm software. Briefly, for brain, regions on interest were drawn to include all brain parenchymal tissue and exclude leptomeningeal areas. For kidney and liver, regions of interest were drawn to include all organ parenchyma. A cell identification and binary categorization algorithm was used to quantify the total number of NuMA-positive cells per slide. Settings for successful discrimination between NuMA-positive and NuMA-negative cells were established for each organ through testing on positive and negative control areas in multiple slides.

### Brain Slice Immunofluorescence

Brains from perfused mice were fixed overnight in 4% PFA. Brains were sectioned using a vibratome (Leica) into 50uM-thick slices. Slices were taken from 3-4 levels evenly spaced through the cortex. Slices were incubated in blocking buffer (10% Normal Goat Serum, 2% BSA, 0.25% Triton) in PBS for 2 hours at room temperature. Primary antibodies were incubated overnight at 4 degrees in blocking buffer and were washed 4 times for 5 min in 0.25% Triton in PBS. Slices were incubated in secondary antibody in blocking solution for 2 hours and were washed 4 times for 5 min in 0.25% Triton in PBS. Nuclei were stained with DAPI at 1:1000 for 4 min in PBS. Brain slices were mounted to glass slides and coverslipped in Dako Fluorescence Mounting Medium (Agilent S3023). Confocal images were captured in z-stack using a Zeiss-770 microscope at 60x in oil. Non-confocal images were captured using a Zeiss-880 microscope at 20x or 40x. The same voltages were used for image capture across all images and groups within each experiment. Images were processed using ImageJ. Within each experiment, brightness and contrast values were kept the same across all images and groups.

The following primary antibodies were used: Anti-GFP AlexaFluor 488 Conjugated Ab (Santa Cruz sc-9996 AF488) 1:200, Anti-Cleaved Caspase-3 AlexaFluor 555 Conjugate (Cell Signaling 9604S) 1:100, Anti-GFAP (Aves Labs) 1:2000, Anti-Iba1 (Wako Chemicals 019-19741) 1:500.

The following secondary antibodies were used: Goat anti-Chicken IgY Alexa Fluor 568 (Thermo Fisher A-11041) 1:500, Goat-anti-Rabbit IgG Alexa Fluor 568 (Thermo Fisher A-11011) 1:500

#### Astrocyte + Microglia Scoring

Confocal (microglia) or non-confocal (astrocyte) images were captured randomly of live cancer cells and their surrounding brain parenchyma. Blinded images were scored according to a qualitative scoring system. For astrocytes, the criteria used to assign a score of 0-4 were brighter GFAP staining than nearby brain parenchyma, extent of astrocyte branching, and extent of GFAP interaction with cancer cells (see Extended Data Figure 4b for example score images). For microglia, the criteria used to assign a score of 0-3 were reactive morphology (less ramified, more ameboid)^42^ compared to nearby brain parenchyma, number of microglia in physical contact with cancer cells, and degree of phagocytosis of cancer cells by microglia (see Extended Data Figure 4c for example score images).

### *Ex-Vivo* Magnetic Resonance Imaging

After perfusion, the skulls of mice were removed and fixed in formalin for 72 hours. MRI experiments were performed on a Biospec 7030 micro-MRI system (Bruker) composed of an Avance-3 HD console and a zero-boil off 7-Tesla (7 T) (300 MHz) 300-mm horizontal bore magnet equipped with an actively shielded gradient coil insert (Bruker BGA-12S-HP; ID 114-mm, 660-mT/m gradient strength, 130-μs rise time). All scans were performed with a Bruker transmit-receive whole mouse body radiofrequency coil (Bruker 1P T20071V3: OD=59mm, ID=38mm, L=40mm) tuned to 300.16 MHz, the ^1^H proton Larmor frequency at 7-T commercial. This rf probe enabled the acquisition of 3D datasets with sub-millimetric isotropic resolution (<150μm) during overnight scans spanning from 8 to 12-hours. As previously described ^22^, tumor burden was detected using multiple sequences. Hyper-intense signal detected by a T_2_-weighted. Rapid Imaging with Refocused Echoes (RARE) sequence recognizes edema surrounding tumors. The 3D RARE sequence was performed with the following acquisition parameters: (120 µm)^3^ isotropic resolution, acquisition time 5h 27 min., repetition time TR = 500 ms, echo spacing ES=12.7 min., Turbo factor TFx=12, effective echo time TE_eff_=76.2 ms, bandwidth BW= 75 KHz, Matrix size = 284^3^, field of view FOV= (4.0 mm)^3^, number of averages Nav=6. Pigmented metastases were detected with signal brightening when using a T_1_-weighted 3D Gradient echo sequence with the parameters as follow: (120 µm)^3^ isotropic resolution, acquisition time 2hrs 41 min., repetition time TR = 20 ms, echo time TE=4.0 ms, flip angle FA= 18°, bandwidth BW= 75 KHz, Matrix size = 284^3^, field of view FOV= (34.0 mm)^3^, number of averages Nav=6. Unpigmented and/or hemorrhagic metastases were detected with a hypo-intense signal when acquired under a T_2_*-weighted multi-gradient echo (MGE) sequence (3D MGE, [120 µm]^3^ isotropic resolution, acquisition time 3h 35 min., repetition time TR = 40 ms, echo time TE=3.6 ms, echo spacing ES=3.2 ms, 4 echoes, flip angle FA= 20°, bandwidth BW= 100 KHz, Matrix size = 284^3^, field of view FOV= (34.0 mm)^3^, number of averages Nav=4. All 3 sequences were used to quantify tumor burden. Identified tumor areas from analysis were cross referenced with histological sections to ensure accuracy.

### Transmission Electron Microscopy

Cultured cells were fixed in 0.1M sodium cacodylate buffer (pH 7.2) containing 2.5% glutaraldehyde and 2% paraformaldehyde for 2 hours and post-fixed with 1% osmium tetroxide for one hour, then block stained in 1% aqueous uranyl acetate, dehydrated using a gradient of ethanol and embedded in EMbed 812 (Electron Microscopy Sciences, Hatfield, PA). Ultrathin sections (60 nm) were cut, mounted on copper grids and stained with uranyl acetate and lead citrate. Stained grids were examined under Philips CM-12 electron microscope and photographed with a Gatan (4k x2.7k) digital camera. Images were analyzed in ImageJ by measuring the largest visible mitochondrial dimension (length or width) for each mitochondrion present in randomly selected images.

### Seahorse Metabolic Analysis

4.5 × 10^4^ 12-273 NBM, 3.5 × 10^4^ 12-273 BM, 8 × 10^4^ 10-230 NBM, 1×10^5^ 10-230 BM, 3 × 10^4^ WM-4071 NBM, or 3 × 10^4^ WM-4071 BM cells were plated on a XF24 Cell Culture Microplate coated with Cell-Tak (Corning). Simultaneously, cells were also plated from same master mix in 12-well plates for later use for normalization. Seahorse MitoStress test protocol was followed using a Seahorse XF24 instrument (Agilent). Concentrations of inhibitors injected are as follows: 12-273 pair – 1uM oligomycin, 1.5 uM FCCP, .5uM antimycin A and rotenone; 10-230 pair .5 uM oligomycin, 1.5uM FCCP, .5uM antimycin A and rotenone; WM-4071 pair -.75 uM oligomycin, 2 uM FCCP, .5 uM antimycin A and rotenone. After MitoStress test, protein was harvested from the normalization 12-well plates using RIPA buffer lysis and quantified by BCA assay. Oxygen consumption rate (OCR) and extracellular acidification rate (ECAR) values for every STC were normalized to average protein concentration of the corresponding normalization wells.

### Plasmid Generation

CMV-Luciferase-EF1α-copGFP (GFP-luc) Lentivector Plasmid was purchased from BD Biosciences (BLIV511PA-1)

#### sh-RNA plasmids

Tet-pLKO-puro was purchased from Addgene (21915). sh-RNAs were cloned as previously described (https://mcmanuslab.ucsf.edu/protocol/cloning-small-hairpins-lentiviral-vectors) into Tet-pLKO-puro using AgeI and EcoRI restriction sites. pLKO tet-on scrambled (sh-Scr) was purchased from Addgene (47541). See Extended Data table 2 for sh-RNA sequences.

#### CRISPR plasmids

pLenti-Cas9 was purchased from Addgene (52962). pLentiGuide-Puro was purchased from Addgene (52963). sg-RNA sequences were designed using the GPP sgRNA Designer (Broad Institute) and cloned into pLentiGuide-Puro using the BsmBI restriction site. See Extended Data table 2 for sg-RNA sequences.

#### APP Expression plasmids

pLVX-IRES-tdTomato (pLVX) was purchased from Clontech (631238). APP-770 was purchased from Genecopoeia (EX-Z2553-M02) and subcloned into pLVX using SpeI and NotI restriction sites.

#### SPA4CT plasmids

A cloning strategy was designed and implemented to generate the SPA4CT sequence ^24^. pLVX APP-770 was digested with EcoRI and NotI to generate a 366 base pair terminal APP insert. Complementary oligos (purchased from Integrated DNA Technologies) were annealed to form an insert with XhoI and EcoRI overhangs. pLVX was digested with XhoI and NotI and religated together with the two inserts. For specific sequences, see Extended Data Table 2. SPA4CT-T43P was generated using the Q5 Site-Directed Mutagenesis Kit (NEB E0554S) using pLVX SPA4CT as a template. For specific sequences of primers and inserts, please see Extended Data Table 2.

### Lentiviral Production and Infection

HEK293T cells at 80% confluency were co-transfected with 12 μg of lentiviral expression constructs, 8 μg of psPAX2 and 4 μg pMD2.G vectors using Lipofectamine 2000 (Invitrogen) following manufacturer’s recommendations. At 48 hr post transfection, supernatants were collected, filtered (0.45 μm) and stored at −80°C. Melanoma cells were infected with lentiviral supernatant supplemented with polybrene at a final concentration of 4 μg/mL. 24 hours after infection, the following selection methods were used for infected cells:

Cell sorting by GFP fluorescence for GFP-luc plasmid.
Culture in puromycin (2ug/mL) for sh-RNA and sg-RNA plasmids.
Culture in blasticidin (10ug/mL) for Cas9 plasmid.
Successful pLVX infection was verified by RFP expression on microscopy; cells were not sorted based on RFP expression.

### In-vitro Proliferation Assay

2 days prior to assay, genetic silencing was induced by addition of doxycycline (1 ug/mL) to media. 1 × 10^4^ 12-273 BM or 1 × 10^4^ WM-4265 BM cells were plated in four 24-well plates. After allowing cells to adhere overnight, a baseline plate was obtained by removal of media and fixation in .1% glutaraldehyde for 15 minutes. Remaining plates were fixed every 24 hours thereafter. Wells were stained with .5% crystal violet in PBS for one hour and washed extensively with water. Crystal violet retained by cells was then eluted by incubation in 250uL 15% acetic acid for 1 hour with shaking. 100uL from each well was transferred to a 96 well plate and absorbance at 590 nm was measured. Absorbances were normalized to the absorbances from the baseline plate to obtain a fold-change value.

### Western Blot Analysis

Cells were harvested in cold RIPA lysis buffer supplemented with protease inhibitor and protein was quantified by BCA assay. Cell lysates (15-20 ug of protein) were resolved in 4%-12% Bis-Tris gels (Invitrogen) and transferred to PVDF membranes using wet transfer. Membranes were blocked in either 5% non-fat milk or 5% BSA in Tris-buffered saline1% Tween 20 (TBST) for 1 hour. Membranes were incubated overnight with primary antibodies diluted in 5% milk or BSA TBST. Membranes were washed 3 times for 10 mins in TBST and incubated in secondary antibody for 1 hour in 5% Milk or BSA TBST. Membranes were washed 3 times for 10 minutes in TBST and incubated with ECL substrate for 3 min and exposed to film for imaging.

The following primary antibodies were used:

Anti-APP 22C11 (Thermo Fisher Scientific 14-9749-82) 1:1000 in milk, Anti-APP C6/1.1 (Gift from Mathews Lab) 1:5000 in BSA, anti-PRKAR2B (Thermo Fisher Scientific PA5-28266) 1:1000 in milk, anti-XPNPEP3 (Atlas Antibodies HPA000527) 1:500 in milk, anti-SCARB1 (Abcam ab52629) 1:1000 in milk, anti-SELENBP1 (Abcam ab90135) 1:1000 milk, anti-EARS2 (Santa Cruz sc-271728) 1:500 in milk, anti-FARP1 (Santa Cruz sc-293249) 1:1000 in milk, anti-MBTD1 (Thermo Fisher 730065) 1:1000 in milk, Anti-Beta-Actin-Peroxidase (Sigma Aldrich A3854) 1:100000 in milk, Anti-tubulin (Sigma T9026) 1:10000 in milk.

The following secondary antibodies were used:

Goat anti-Rabbit IgG-Peroxidase (Sigma-Aldrich A0545) 1:5000, Goat anti-mouse IgG kappa-light chain (Santa Cruz sc-516102) 1:1000.

### Membrane purification and γ-secretase activity assay

Cell membrane preparation and γ-secretase assays were described previously ^75–78^. Briefly, cells were harvested from T75 flask 90% confluency and collected by centrifuge (800 g, 10 min). Cell pellets were resuspended with hypotonic buffer (5mM Tris, pH 7.4), incubated in ice for 30 min and homogenized with Glass Teflon Homogenizer. Samples were centrifuged at 1000 g, 30 min and the supernatants that contain the total membrane fraction were collected. Pellets were resuspended with hypotonic buffer, homogenized and centrifuged again. Combined supernatants were centrifugated at 100,000g for 60 min. Resulted pellets referred to as membrane fractions were resuspended and washed with MES buffer (50 mM MES, pH 6.0, 150 mM KCl, 5 mM CaCl2, 5 mM MgCl2, and protease inhibitors) and spun down (100,000g for 60 min). Membrane fractions were dissolved in MES buffer and protein concentration was determined by the DC assay kit (Biorad). For γ-secretase assays, Sb4 substrate (1 µM) or NTM2 substrate (0.4 µM) and membrane fractions (50 µg/ml) were incubated in PIPES Buffer (50 mM PIPES, pH 7.0, 150 mM KCl, 5 mM CaCl2, 5 mM MgCl2) and 0.25% CHAPSO detergent at 37°C for 3 hours. *γ*-Secretase products were detected by AlphaLISA methods using G2-10 or SM320 antibodies for Aβ40 or NICD, respectively.

### Amyloid Beta ELISA

Media was conditioned with melanoma cells for 24-72 hours and concentrated 10x using Amicon Ultra 3-kDa Concentrators (Millipore Z740169). Amyloid Beta-40 ELISA was performed using the Human AB40 ELISA Kit (Invitrogen). Secretion values were normalized to protein content of wells as measured by RIPA harvest and BCA protein quantification.

### APP Isoform Semi Quantitative qPCR

RNA was isolated from 10-230 NBM, 10-230 BM, 12-273 NBM, and 12-273 BM cells at 80% confluence in 6 well plates using the RNeasy Mini Kit (Qiagen 74104). 600 ng of RNA was subjected to DNase I treatment and reverse transcription. PCR was performed with primers as previously described ^79^to generate bands of different sizes corresponding to the following isoforms: APP-770 – 461bp, APP-751 – 417bp, APP-695 235bp

### Statistical Analysis

Statistical analyses were performed with Prism 8 (GraphPad Software). Unless otherwise stated, the Student’s t-test was used for experiments. P-values <0.05 were considered to be statistically significant. Unless otherwise stated, values are averages and error bars are +/− standard deviation.

## Acknowledgements

E.H. is supported by NIH R01CA2022027, P01CA206980, a Leveraged Finance Fights Melanoma-MRA Team Science Award and a NIH Melanoma SPORE (NCI P50 CA016087; PI: I.O.). K.K. is supported by F30CA221068, and previously by the NIH/NCI 5 T32 CA009161-37 (Training Program in Molecular Oncology and Immunology, PI: D.E. Levy), NIGMS 5 T32 GM007308-41 (Medical Scientist Training Program, PI: M. Philips), and a Vilcek Foundation Scholarship. A.F. is supported by a Fundacion Ramon Areces fellowship. YM.L. is supported by RF1AG057593. S.L is funded by Cure Alzheimer’s Fund, Anonymous Donors and the NYU School of Medicine.

We thank Drs. Clemens Krepler and Meenhard Herlyn for patient-derived short-term cultures. We thank Dr. Rana Moubarak and members of the Hernando lab for critical reading of the manuscript. We thank the NYU Center for Biospecimen Research and Development (CBRD), the Experimental Pathology Core Facility (Director, Dr. Cindy Loomis), the Immunohistochemistry Core Laboratory (Dr. Luis Chiriboga), the Microscopy core (Dr. Alice Xiang), the Genomics Technology Center (Dr. Adriana Heguy), and the Flow Cytometry Core (Dr. Peter Lopez), supported in part by the Laura and Isaac Perlmutter Cancer Center Support Grant NIH/NCI P30CA016087, and National Institutes of Health S10 Grants NIH/ORIP S10OD01058 and S10OD018338.

## Author Contributions

K.K., R.J.S. and E.H conceived and designed the experiments. K.K, assisted by G.L., performed the experiments and analyzed the data. G.L. performed *in-vitro* and *in-vivo* experiments with WM-4265 BM. E.W. conducted gamma-secretase *in-vitro* assays, supervised by Y.L. F.G-E. assisted with generation of mutant APP constructs and Aβ ELISA analysis. R.V-I. and D.A. assisted with in-vivo experiments. I.R. isolated astrocytes, supervised by S.L. L.B. performed bioinformatics analysis of RNA-seq data, supervised by K.R. A.F. performed pathway analysis of proteomics data and western blot analysis of STC pairs. J.R. and J.C. analyzed ex-vivo MRI data, supervised by Y.Z-W. A.D. performed mass spectrometry analysis, supervised by B.U. E. deM. and I.O. generated and provided patient-derived STCs. R.R performed histological analysis of in-vivo experiments using paired STCs. P.M. assisted with design of Aβ immunoprecipitation from melanoma conditioned media. K.R. performed statistical and pathway analysis of proteomics data. S.L. assisted with design and analysis of *in-vitro* astrocyte experiments. R.D. provided BACE-i and assisted with BACE-i experimental design. K.K and E.H. wrote the manuscript.

